# Interpreting Sleep Activity Through Neural Contrastive Learning

**DOI:** 10.1101/2024.09.25.615100

**Authors:** Zhongtao Chen, Hui Zheng, Jianyang Zhou, Lin Zheng, Peiyang Lin, Haiteng Wang, Marc Busche, Tim Behrens, Ray Dolan, Yunzhe Liu

## Abstract

Memories are spontaneously replayed during sleep, a process thought to support memory consolidation. However, capturing this replay in humans has been challenging because unlike wakefulness, sleep EEG is dominated by slow, rhythmic background activity. Moreover, each sleep stage (e.g., NREM, REM) has distinct rhythms, hindering generalisation of models trained on wake-state data. To overcome these challenges, we developed the Sleep Interpreter (SI), a neural network model that decodes memory replay from sleep EEG. In a large dataset comprising 135 participants (∼1,000 h of overnight sleep; ∼400 h of wake), we employed a TMR-like paradigm with 15 semantically congruent cue-image pairs to tag specific memories. SI was trained separately for NREM and REM using contrastive learning to align neural patterns across wake and sleep, filtering out stage-specific background rhythms. We also examined how slow oscillations and spindle coupling influence decoding in NREM sleep. In a 15-way classification, SI achieved up to 40.02% Top-1 accuracy on unseen subjects. To test generalisability, we followed up with two independent nap experiments in separate samples and applied the trained SI model off-the-shelf. The first probed spontaneous reactivation without auditory cues, while the second used semantic-free sounds with new images. In both, SI successfully decoded reactivation during sleep that correlated with post-nap memory performance. By openly sharing our dataset and SI system, we provide a unique resource for advancing research on memory and learning during sleep, and related disorders.

## Introduction

Sleep is considered essential to cognition^1,2^. In particular, it provides a state in which the brain spontaneously replays prior experiences, a process thought to underlie memory consolidation^3–5^. This replay is particularly active during non-rapid eye movement (NREM) sleep, where the brain shifts into slower, more synchronised activity patterns^6,7^. While memory replay during sleep has been widely demonstrated in animal models^8,9^, capturing this in humans remains a significant challenge.

Human sleep activity, as typically measured in EEG, is dominated by rhythmic patterns, including large-amplitude (>100 µV), low-frequency slow oscillations (SOs) seen in deep, N3-stage sleep^10,11^, which swamp the few-microvolt signals associated with memory replay. Past attempts to decode sleep content often required extensive feature engineering and off-line analyses^12,13^. However, bridging this “generalisation gap” between wake and sleep data demands methods capable of extracting subtle cross-domain representations despite vast differences in temporal scale and signal-to-noise ratio^14,15^.

This challenge is further exacerbated by recording constraints: in humans, only non-invasive methods such as EEG, MEG or fMRI are typically available. In animal studies, researchers can directly record neuronal spikes, which exhibit similar representations across sleep and wake^16,17^. Human EEG and other non-invasive recordings cannot capture such fine-grained neuronal activity owing to limited temporal and spatial resolution. Additionally, stark differences in background activity and target signals between wakefulness and sleep, along with difficulties in aligning relevant time scales, further widen the generalisation gap.

Decoding neural activity using machine learning provides a powerful way to link brain states with specific perceptual or cognitive content^18^. Approaches range from linear classifiers (e.g., LDA, SVM)^19–22^, to deep-learning models (e.g., EEGNet^23^). In awake individuals, EEG decoders can distinguish visual categories^24–27^, semantic classes^28–31^, and motor intentions^23,32–35^, often with accuracy exceeding 70% in four-class tasks^35–38^. However, extending such decoding approaches to sleep has proved challenging^18^.

In this study, we tackle these challenges by developing a sleep-decoding system trained on a large dataset specifically acquired to enable sleep decoding. Our dataset comprises recordings of whole-brain EEG activity from 135 participants during both wakeful tasks and overnight sleep (∼1 000 h of sleep; ∼400 h of wake). Using a simplified, learning-free Targeted Memory Reactivation (TMR)-like paradigm^39–42^, we paired 15 semantically congruent auditory cues with visual images during wakefulness and replayed these cues during sleep to track memory reactivation events. This design enabled us to train the Sleep Interpreter (SI) model on both wake and sleep reactivations.

To align representations of identical semantic content across wakefulness and sleep, we employ neural contrastive learning^43^ operating in a sleep-stage-specific manner to account for background activity differences between stages^44,45^. Unlike conventional contrastive approaches applied to generic feature spaces^43,46^, our method aligns raw neural signals of identical semantic items across states. By linking reactivation patterns in sleep to their corresponding wake representations—thereby filtering out stage-specific background rhythms such as dominant SOs in NREM sleep^10,11^—SI isolates semantic-related signals and mitigates interference from background activity.

NREM sleep is characterised by SO-spindle coupling and is thought to be the primary window for memory replay^39,47–49^, whereas REM sleep displays a desynchronised, wake-like EEG associated with dreaming^50,51^. To accommodate these differences, we trained separate SI models for NREM and REM and developed an online sleep-staging module to switch between them in real time. Focusing on stage-relevant features, such as SO-spindle coupling in NREM, our real-time system can track memory reactivations throughout the night.

To further test SI’s generalisability outside our initial paradigm, we conducted two independent nap experiments inspired by prior sleep research^6,39,40,52–55^, each with a separate participant sample, and applied the pre-trained SI model off-the-shelf to decode memory reactivation during sleep. The first experiment (n=25) probed spontaneous memory reactivation without any auditory cues^39^, while the second experiment (n=22) used a classic TMR paradigm with semantic-free sounds and novel images^6^. In both cases, SI successfully decoded reactivations that correlated with post-nap memory performance, demonstrating its applicability across different tasks, materials, and samples, and indicating that SI performance is not driven by specific properties of the eliciting auditory cues.

## Results

### Task design and dataset

Establishing effective real-time sleep decoding first necessitates **ground truth** data that indexes *which* memories get reactivated and *when*. For this we developed a variant of a TMR task^40^ (TMR-like task) with simultaneous 64-channel scalp EEG recordings acquired during both awake tasks and overnight sleep (**Figure 1a**). Data were recorded at 500 Hz from 135 healthy, university-educated adults (mean age 22.55±0.22 years; 72 female). In total, the dataset comprised 957.03 hours of sleep and 362.29 hours of wake recordings (see details in **Methods**).

**Figure 1.**
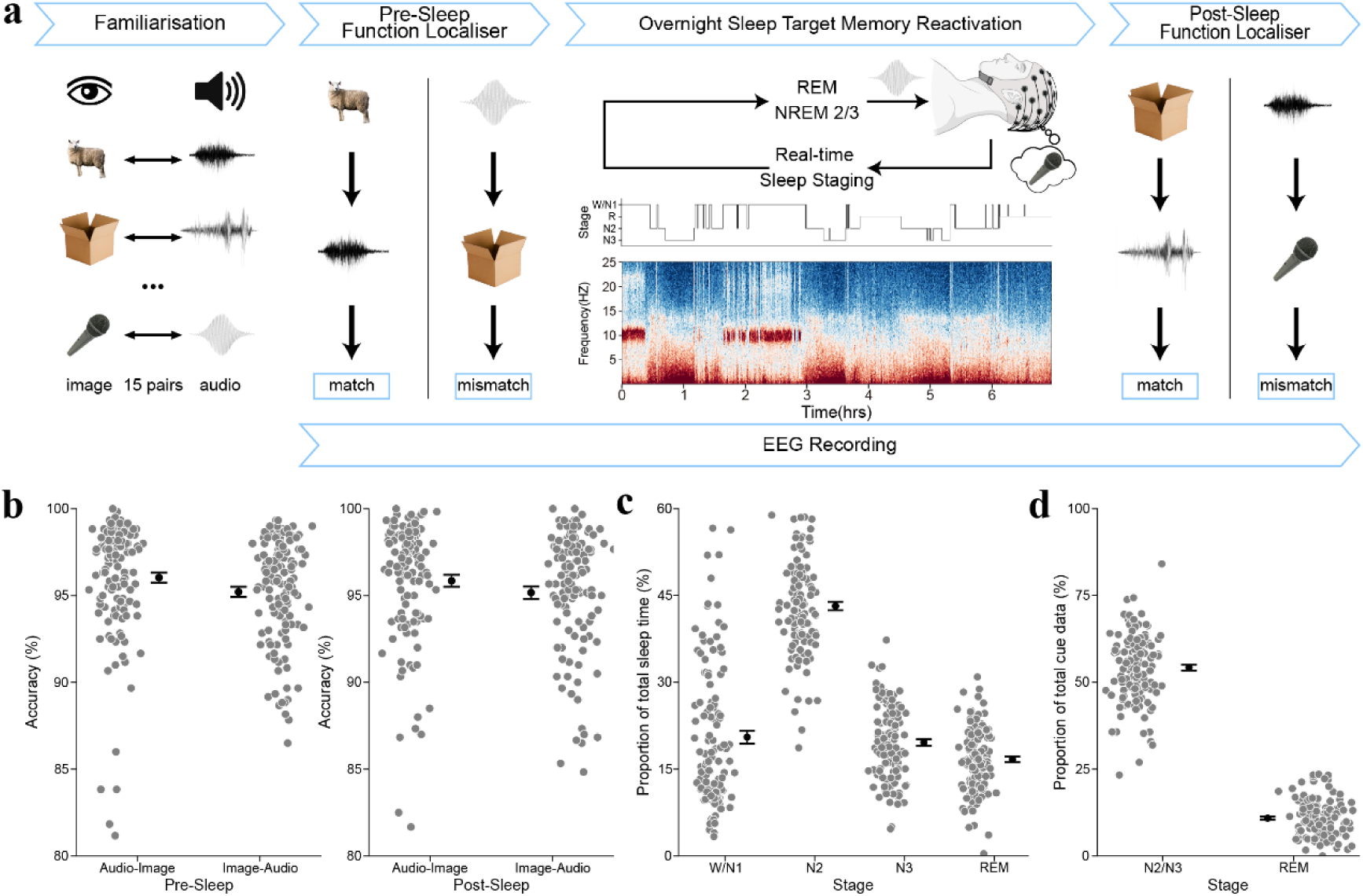
The experimental design for sleep decoding. (**a**) The experiment began with a familiarisation stage wherein subjects learned 15 predefined, semantically congruent picture-sound pairs (e.g., a picture of a “sheep” was paired with the sound of a sheep ‘baa’). Following this familiarisation stage, subjects underwent a functional localiser task involving the presentation of 1200 trials in total. Each trial contained one picture and one sound selected from the 15 learned picture-sound sets (match or mismatch). These were presented in either image→ audio or audio→ image order, and subjects were asked to determine whether the pairs were matched or not. Subsequently, during overnight EEG recording, real-time online sleep staging was performed. When subjects entered NREM 2/3 or REM sleep stage, auditory cues—randomly selected from the image-audio pairs—were played every 4-6 seconds. This step provided an approximation to a ground truth regarding which picture memory was likely to be activated and when. Following overnight sleep, subjects were again presented with the 1200 image-audio pairs. Whole-brain 64-channel EEG recordings were collected throughout the experiment, during both sleep and functional localiser tasks. More details can be found in **Methods**. (**b**) During pre- and post-sleep functional localiser stages, subjects were tasked to determine whether the presented image and audio cues were congruent. Behavioural accuracy is shown separately for image→audio and audio→image, as well as for pre-sleep and post-sleep periods. (**c**) The proportions of different sleep stages as a function of total sleep time, as determined by the offline, expert-based manual sleep staging. (**d**) The proportions of cued sleep data in N2/3 and REM, following offline, expert-based manual sleep staging. The gray dots represent the results for each subject, while the black error bars indicate the mean (M) and standard error of the mean (SE) for the corresponding conditions.

During wakefulness, subjects were first familiarised with 15 semantically congruent sound-image pairs, each representing a distinct concept (e.g., ‘sheep’, ‘book’). Only after subjects confirmed knowledge of the correct pairings did they proceed to a functional localiser phase. In the latter, each trial presented one image and one sound in random order, while participants judged whether the semantic content of the pair matched. This localiser session was conducted twice: once before sleep (pre-sleep) and again following overnight sleep (post-sleep). In the picture-sound pairing functional localiser sessions test subjects performed well behaviorally (**Figure 1b**), both pre-(image → audio, 95.21±0.30%; audio → image, 96.04±0.29%) and post-sleep (image → audio, 95.17±0.35%; audio → image, 95.85±0.35%), with no significant difference between them (image→audio: *t*(134)=0.21, *P*=0.836; audio→image: *t*(134)=0.84, *P*=0.403, two-sided paired *t*-test).

Only correctly responded trials with good EEG data quality were epoched (from 0 to 0.8 seconds following the onset of the image or audio stimulus) for later neural decoding analysis (see details in **Methods**). Across subjects, for both image- and audio-evoked EEG, signals were segmented into 2153.42±29.08 sample pairs data. Each sample was paired with a corresponding label from the 15 predetermined semantic concepts. For an identical semantic concept, neural data evoked by either images or audio clips during wakefulness were later used to align with audio-induced neural representations elicited during sleep.

The mean sleep time across subjects was 7.09±0.06 hours (measured across getting into bed to waking up in the morning; **Extended Data Table 1**). During actual sleep we monitored sleep stages in real-time including once subjects entered NREM 2/3 or REM sleep (determined by an experienced experimenter). Each audio cue, randomly chosen from the 15 pairs, was presented at 43.51±0.38 dB, with 4-6 s intervals between successive clips to prevent overlap of conceptual representations. In offline analysis, for each subject, two sleep experts independently annotated the sleep stage (Wake, N1, N2, N3, and REM) in 30-second segments covering the entire night. In cases of inconsistency, the experts engaged in a discussion to reach a consensus. On average, subjects spent 20.53±1.10% of their sleep time in Wake and N1 stages, 43.18±0.72% in N2 stage, 19.61±0.57% in N3 stage, and 16.68±0.50% in REM stage (**Figure 1c**). In relation to auditory cues delivered during the night, 34.88±0.87% were during the W/N1 stage, 54.25±0.84% during NREM 2/3 stage, and 10.87±0.43% during REM stage (**Figure 1d**). After excluding bad segments and auditory cues presented in the W/N1 stage, nocturnal TMR-related EEG signals were segmented into 1383.30±40.97 samples for NREM 2/3 stage, and 307.66±15.48 samples for REM stage. Each sample was epoched from -0.2 to 1.8 seconds after the onset of the audio stimulus and paired with the corresponding semantic label.

In total, we obtained 290,712 pairs of image- and audio-evoked EEG samples from the awake tasks (pre- and post-sleep combined), and 186,746 NREM 2/3 and 41,534 REM audio-evoked EEG samples during sleep (**Extended Data Table 2**). These samples were annotated and curated into a dataset for sleep decoding. To the best of our knowledge, this is the first dataset that permits training neural decoders for the analysis of sleep content. The dataset is accessible from https://osf.io/x7453/ upon publication.

After dataset curation, we developed a sleep decoding system in three key steps. First, we built an automatic sleep staging model to classify sleep stages without the need for manual intervention. Next, using contrastive learning to align neural representations between wakefulness and sleep, we designed a sleep content decoder. Finally, we incorporated sleep rhythms associated with memory consolidation, selecting specific features to further improve the model’s decoding accuracy. The final zero-shot accuracy on unseen subjects reached up to 40.02% for 15-class classification during NREM sleep, outperforming all available EEG decoding models. The sleep decoding system was then tested in two new experiments, verifying its capacity to detect memory reactivation and to generalise across diverse experimental conditions. We term our model ‘Sleep Interpreter’, or SI, which in Chinese pronunciation is ‘思’, meaning ‘thought’.

### Automatic real-time sleep staging model based on ANN

Sleep staging is typically carried out offline via a human agent using a visual scoring protocol^56,57^. Although automated sleep staging algorithms, such as YASA^58^, are available, their real-time capabilities are undetermined. A real-time capability is crucial for controlling timing of cued memory reactivation during sleep, including during distinct sleep states.

As we are interested in zero-shot generalisation across subjects, we used expert-labelled sleep staging as a ground truth to train a real-time sleep staging model using an artificial neural network (ANN) (**Figure 2a**; see details in **Methods**). The model was trained on sleep EEG data from three channels (C3, EOG, EMG) across 123 subjects and tested on sleep data from the remaining 12 subjects (randomly chosen, roughly 10% of the training data). An example of a sleep staging result on testing subjects is shown in **Figure 2b**, alongside its ground truth.

**Figure 2.**
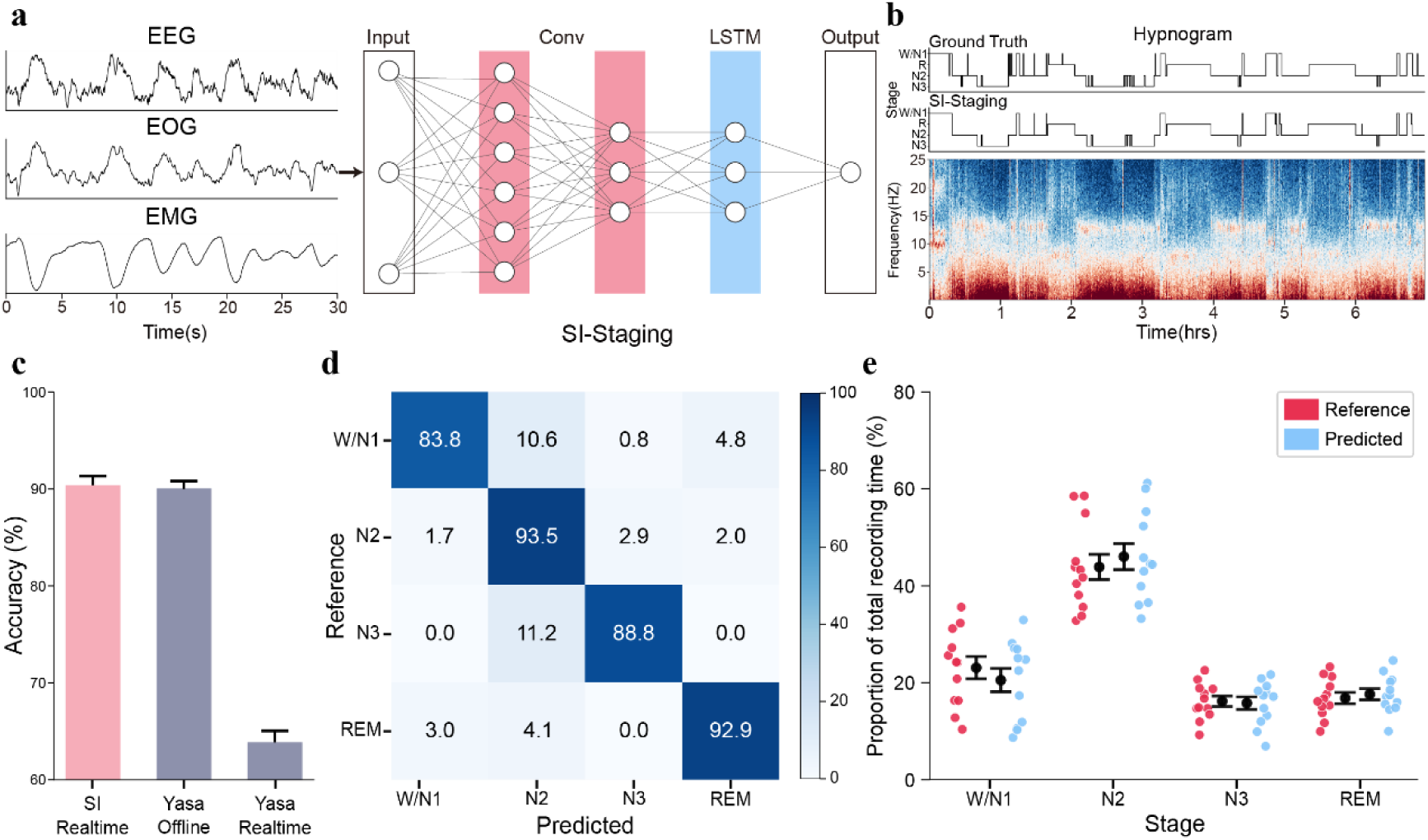
Real-time sleep staging. (**a**) Our Sleep Interpreter (SI) staging model architecture comprises a CNN block, an LSTM network and a classification head. To maximise staging accuracy, the model uses 30-second data segments from three channels (EEG-C3, EOG, EMG) as inputs, filtered in a frequency band consistent with YASA^58^. The model’s loss function incorporates both cross-entropy and mean-square error terms, a combination that aims to address an imbalance in the distribution of data volume across each sleep stage, thereby enhancing the accuracy of sleep staging. (**b**) Ground-truth (expert-labeled) and model real-time predicted hypnogram for one test subject. The agreement between the ground-truth and model real-time predicted scoring is 94.41%. Below the hypnogram is the corresponding full-night temporal frequency profile of the central electroencephalogram (EEG) channel – C3, with warmer colours indicating higher power in these frequencies. (**c**) The mean consistency with the ground truth (expert-labeled) for different staging models. Our real-time SI-Staging model is represented in light pink, while the two YASA-based models are shown in dark blue. For the YASA real-time model, calculations are based on the past five minutes of data due to the model’s limitations. (**d**) Confusion matrix of staging accuracies. The diagonal elements represent the percentage of epochs correctly classified by the SI-Staging model, whereas the off-diagonal elements show the percentage of epochs mislabeled by the SI-Staging model. (**e**) Proportion of each sleep stage in the total sleep time: ground truth (reference, in red) vs. the SI-Staging model predicted (in blue). The dots represent the results for each subject, while the error bars indicate the mean (M) and standard error of the mean (SE) for the corresponding conditions.

Across 12 test subjects our SI-Staging model achieved an average consistency with a ground truth of 90.38±0.96%. This was numerically higher, though not statistically significant, compared to the 90.05±0.76% consistency achieved by YASA^58^, an offline staging algorithm (*t*(11)=0.40, *P*=0.700, two-sided paired *t*-test; **Figure 2c**). When YASA^58^ was used in real-time, its consistency with a ground truth was only 63.86±1.19%, with our real-time staging model performing significantly better (*t*(11)=14.55, *P*<0.001, two-sided paired *t*-test).

In classifying each sleep stage our staging model also showed good specificity (**Figure 2d**). The mean consistency with a ground truth was 88.8±4.05% for the N3 stage, 93.5±1.24% for the N2 stage, and 92.9±1.50% for REM sleep stage. For wakefulness and N1 stage, categorised together, its consistency was 83.8±3.07%. Similarly, the model predicted time for each sleep stage was comparable with the ground truth (expert-labelled) sleep duration (**Figure 2e**). The proportion of N2 or N3 stage in the total sleep time was 43.89±2.61% or 16.19±1.09%, with no significant difference compared to the model prediction (*t*(11)=1.24, *P*=0.228, two-sided paired *t*-test). The same was true for REM stage (16.83±1.17%, *t*(11)=1.33, *P*=0.211, two-sided paired *t*-test). These results demonstrate that, when compared to offline expert-labelled stage results, our SI-Staging model achieves both good consistency and specificity in sleep staging.

### Generalisation ability of neural representations across awake and sleep states

To decode reactivation of objects during sleep, we require access to their neural representations, which are typically derived from training during an awake task. Sleep decoding therefore relies on the assumption that object representations generalise across awake and sleep domains. In the present task, we could compare event-related potentials (ERPs) elicited by the same semantic object during awake and sleep states. Qualitatively, there is a striking domain difference between awake and sleep ERPs (**Figure 3a**): sleep ERPs (audio-evoked) are much more synchronised across channels than both image- and audio-evoked ERPs obtained during the awake task (for ERP differences between NREM and REM sleep, see also **Extended Data Figure 1**). There is also a discernible difference between audio-evoked and image-evoked ERPs, although to a lesser degree.

**Figure 3.**
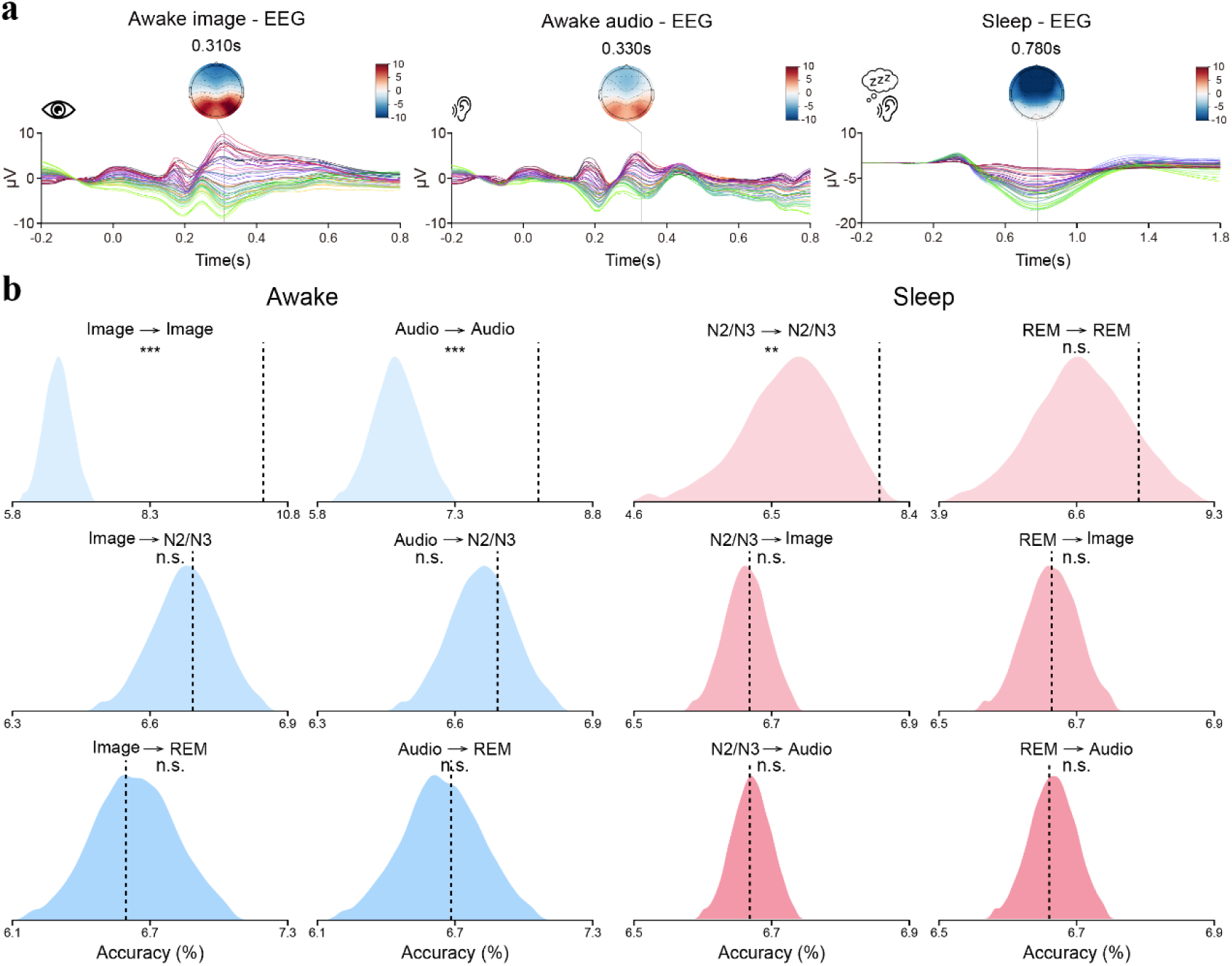
ERPs in awake and sleep domains, their within-domain decodability, and across-domain generalisation ability. (**a**) Averaged ERPs and topographic maps evoked by image stimuli during wakefulness, audio stimuli during wakefulness, and TMR-related audio stimuli during NREM-2/3 and REM sleep, respectively. Zero on the horizontal axis marks stimulus onset for each domain. Awake ERPs were averaged from −0.2 s to +0.8 s relative to onset; sleep ERPs were averaged from −0.2 s to +1.8 s. Topographic maps display time points with the strongest ERP responses. (**b**) Permutation-based null distributions (1,000 label shuffles) of decoding accuracy, for cross-domain decoding (awake → sleep, blue densities; sleep → awake, red densities) and within-domain decoding (same train and test state; awake in blue, sleep in red). The classifier is a Lasso-regularised GLM. Black dashed lines indicate decoding accuracy on unpermuted, real data. *n.s.*, non-signifciant, ** *p*<0.01, *** *p*<0.001.

With these considerations in mind, we first tested within-domain decodability and cross-domain generalisability using a Lasso-regularised GLM, whose L1 penalty imposes a non-linear sparsity constraint for effective feature selection^59–62^ (**Figure 3b**). For each subject, we trained object decoders on image-evoked (−0.2 s to 0.8 s from stimulus onset) or audio-evoked (−0.2 s to 0.8 s) ERPs in the awake state and tested their performance both within the awake state and across sleep states. Similarly, we trained decoders on audio-evoked ERPs acquired during sleep (−0.2 s to 1.8 s) and tested them in both sleep and awake states. Note that within this TMR-like paradigm we know which object is likely to be activated and when. We implemented this approach separately for NREM and REM sleep, with time windows chosen to match the later SI model training, itself defined through an initial search for optimal sleep-decoding accuracy (see **Methods** for details).

The model was trained and tested within each subject using data from either identical domains (e.g., awake image vs. awake image) or from different domains (e.g., sleep vs. awake). For within-domain decoding, we applied a 5-fold cross-validation procedure and averaged the resulting performance metrics. Conversely, cross-domain analysis was conducted by directly training and testing without cross-validation. To statistically assess the decoding feasibility in both analyses, we employed permutation testing, repeating identical pipelines 1,000 times with shuffled training labels. This procedure generated a null distribution against which the actual decoding results were compared.

Within-domain decoding using the awake image-based decoder yielded an accuracy of 10.54±0.14 %, whereas the awake audio-based decoder yielded 8.21±0.06 %. Both results exceeded their respective 95th-percentile permutation thresholds, demonstrating performance significantly above chance (*P*<0.001, permutation test). However, generalisation from these awake-trained decoders to NREM-2/3 sleep produced accuracies of 6.69±0.01 % for both decoder types, which were not significantly higher than chance (image-based: *P*=0.443; audio-based: *P*=0.317). Decoding accuracies during REM sleep were similarly low: 6.59±0.03 % and 6.68±0.03 % for awake image- and audio-based decoders, respectively, neither significantly surpassing chance levels (image-based: *P*=0.581; audio-based: *P*=0.409; three subjects excluded owing to no REM sleep data).

Conversely, within-doamin decoding performance neural decoders trained and tested within NREM and REM sleep states reached accuracies of 7.99±0.07% and 7.47±0.07%, respectively, the NREM result was significantly above chance, whereas the REM result was not (NREM: *P*=0.003; REM: *P*=0.123). Nevertheless, generalisation from NREM sleep-trained decoders to awake state data yielded accuracies of 6.67±0.01% for both awake-image and awake-audio conditions, not significantly exceeding chance performance (awake-image: *P*=0.436; awake-audio: *P*=0.531). Similarly, REM sleep-trained decoders tested on awake data achieved accuracies of 6.66±0.01% for both awake-image and awake-audio conditions, again neither significantly surpassing chance (awake-image: *P*=0.493; awake-audio: *P*=0.564).

Together, these findings show that while within-domain decoding was robust and consistently surpassed permutation-based thresholds, none of the cross-domain accuracies exceeded the critical 95th-percentile criterion. We therefore conclude that, although the Lasso GLM captures stimulus-specific neural information within a given state, it is inadequate for decoding congruent representations across wakefulness and sleep. Consequently, we turned to a deep-learning model capable of capturing richer, non-linear neural representations to enhance cross-domain generalisation.

### Decoding object representation during sleep with neural contrastive learning

We next developed a cross-domain, neural-network-based decoding model, the SI-Multi-Domain (SI-MD). This model employs neural contrastive learning^43^, to explicitly align each object representation during wakefulness with its counterpart during sleep (**Figure 4a**; see details in **Methods** and **Extended Data Table 3**). The critical computations of the SI-MD model involve two parts: the model architecture and the inclusion of neural contrastive learning loss to align representations across awake and sleep domains.

**Figure 4.**
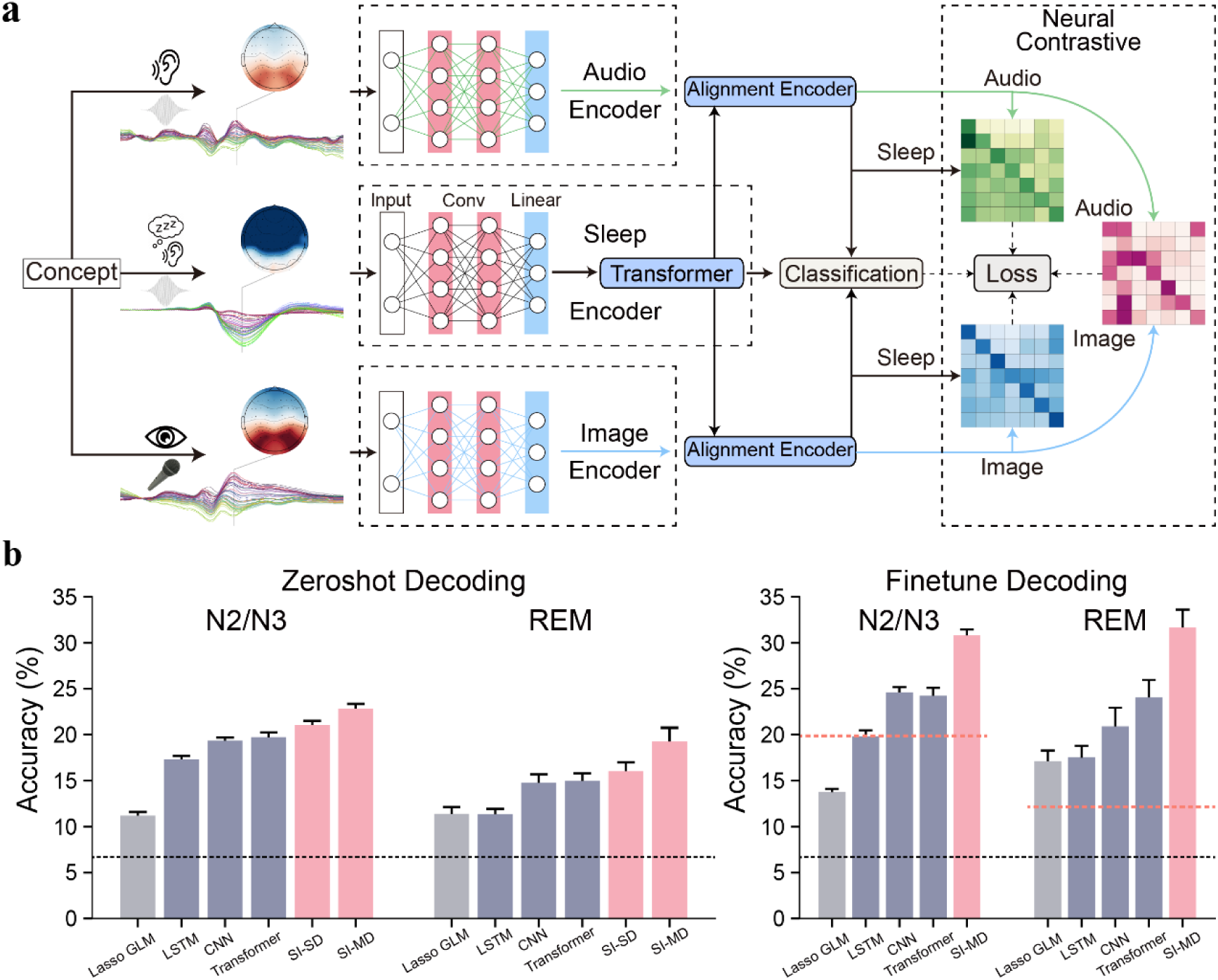
Zero-shot cross-subject sleep decoding with neural contrastive learning. (**a**) Overview of Sleep Interpreter – Multi Domain (SI-MD) model. The SI-MD model architecture consists of neural encoders for both sleep and awake datasets, as well as the alignment encoders. Contrastive loss is used to regularise the latent space, encouraging similarity among features adhering to the same semantic class but coming from different domains (e.g. 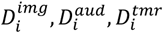) of the same subject, *i*. The awake EEG dataset and sleep EEG dataset are perceived as disparate domains due to the substantial differences in their neural patterns. By utilizing the alignment encoder approach and integrating neural contrastive learning, the model effectively aligns awake and sleep neural representations across subjects during training. (**b**) Sleep decoding performance. The zero-shot, cross-subject decoding performance of all tested models is shown on the left, and the performance of fine-tuned models is shown on the right, separately for the NREM 2/3 stage and REM sleep stages. The Lasso GLM model is represented by gray bars, widely used ANN models by dark blue bars, and the SI-SD and SI-MD models by light pink bars. The black dashed line represents the chance level accuracy for the decoding task, which is 1/15. The orange dashed lines represent the average result of the SI-SD model trained and tested on individual subjects across the twelve test subjects, which is 19.87±0.88% for the NREM 2/3 stage and 12.13±1.01% for the REM stage. The bar charts display the mean performance (M) with error bars representing the standard error of the mean (SE) across the testing subjects.

The model architecture of SI-MD consists of three encoders, one for each data domain: ‘awake image’, ‘awake audio’, and ‘sleep audio’ (**Figure 4a**). For each encoder, the stimuli-evoked whole-brain EEG activity is passed through a one-dimensional (1D) convolution module^63^ and a linear module. Additionally, to capture global synchronised brain activation patterns during sleep, the sleep encoder goes through a transformer module^64^. Finally, to decode the identity of an object, each processed data stream goes through a classification layer. Each pair of data streams also share an encoder for calculating neural contrastive loss (see details in **Methods**).

The typical contrastive learning loss is calculated as:

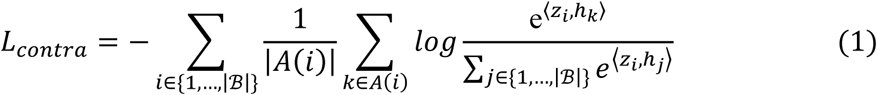

where *A*(*i*) = {*k*|*k* ∈ {1,…, |ℬ|}, *y*_*k*_ = *y*_*i*_}, ⟨⋅,⋅⟩ is the inner product, B is the batch dataset with |ℬ| the size of batches and y represents the semantic label. Typically, *z* and ℎ represent data from different domains. In neural decoding^65^, *z* could represent latent embeddings from neural activity, while ℎ could represent processed sound from the wav2vec model^66^.

By changing *z* and ℎ to represent latent embeddings from different kinds of EEG signals (e.g., sleep-TMR-related and image-evoked EEG signals), we derived a neural contrastive learning loss function.

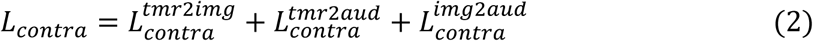

In essence, this method aligns the neural representation of the same object across domains, facilitating sleep decoding, and where our expectation was better decoding accuracy compared to models trained on sleep data alone. We then went on to compare the performance of our SI-MD model with other ‘control’ models. All sleep decoding models (SI-MD and other control models) were trained on 123 subjects and tested on the remaining 12 subjects. This was implemented separately for NREM 2/3 and REM sleep stage. Note we are particularly interested in the zero-shot, cross-subject, generalisation ability of the sleep decoder, given our aim is to derive a model that can decode sleep in subjects outside the framework of current study.

To evaluate the impact of neural contrastive learning loss, we compared the performance of the SI-MD model with a model that shares the same architecture but lacks this cross-domain contrastive learning, and only trained on the sleep data domain, termed SI-Single Domain (SI-SD). The SI-MD model achieved zero-shot accuracies of 22.82±0.52% and 19.27±1.47% at NREM and REM stages respectively, significantly higher than the SI-SD model’s zero-shot accuracies of 21.07±0.43% and 16.05±1.59% (**Figure 4b**, NREM2/3: *t*(11)=4.61, *P*<0.001; REM: *t*(11)=3.19, *P*=0.008, two-sided paired *t*-test). Furthermore, after fine-tuning, the decoding accuracy of the SI-MD model during sleep for the same subjects improved to 30.81±0.65% for NREM 2/3 stage and 31.65±1.96% for REM stage.

To evaluate model architectures, we implemented five different types of sleep decoding models: our SI-MD model, SI-SD model, a linear decoder model (Lasso GLM), and three ANN models (LSTM-based, CNN-based, and Transformer-based). Details on model construction are available in the ‘Decoding model comparison and ablation experiment’ section in **Methods**.

We found that the zero-shot decoding results for all models significantly surpassed chance level (**Figure 4b**, NREM2/3: all *t*(11) > 11, all *P*<0.001; REM: all *t*(11) > 6.30, all *P*<0.001, two-sided one-sample *t*-test). After fine-tuning on sleep task data, the decoding performance of our SI-MD model, as well as the CNN-based and Transformer-based models, significantly exceeded those trained on sleep data alone (NREM2/3: all *t*(11) > 3.30, all *P*<0.01; REM: all *t*(11) > 3.50, all *P*<0.01, two-sided paired *t*-test), whereas the Lasso GLM model performed significantly worse (*t*(11) = - 7.47, *P*<0.001, two-sided paired *t*-test). Moreover, our SI-MD model significantly outperformed the other four models in terms of both zero-shot accuracy (NREM2/3: all *t*(11) > 8, all *P*<0.001; REM: all *t*(11) > 4, all *P*<0.01, two-sided paired *t*-test) and fine-tuning accuracy (NREM2/3: all *t*(11) > 11, all *P*<0.001; REM: all *t*(11) > 6.30, all *P*<0.001, two-sided paired *t*-test). This indicates wide applicability of our SI-MD model, including the potential to integrate awake task data for decoding sleep in unseen subjects.

We also compared the performance of SI-MD model to widely used decoding models, such as EEG-Net^23^, EEG-Conformer^67^, BrainBERT^68^, BIOT^69^ and LaBram^70^. None of those models achieved zero-shot, cross-subject, decoding accuracy above 20% when applied to NREM 2/3 stage and 15% in REM stage (**Extended Data Table 4**). Further control analysis revealed that sufficient data benefits training the SI-MD model, with decoding accuracy increasing as the training data size increases (**Extended Data Figure 2**). Training on the broadband data signal was more effective than training on specific frequency bands (i.e., delta, theta, alpha, beta, gamma) for both NREM 2/3 and REM sleep stages (**Extended Data Figure 3**).

Together, these results suggest that our SI-MD model, incorporating neural contrastive learning that aligns neural representations across awake and sleep domains, outperforms an SI-SD model trained on sleep data alone. Crucially, it also outperforms other widely used ANN architectures and EEG-based decoding models, a performance advantage that held for both NREM 2/3 and REM sleep stages.

### Sleep decoding performance as a function of cue time in SO phases

Previous studies show that, during NREM sleep, the strength of spontaneous memory reactivation varies across different phases of a Slow Oscillation (SO) event^71^. Reactivation is stronger in the “UP” state, defined as the peak of the SO, than in the “DOWN” state, defined as the trough of the SO^6,55^. There is also coupling between spindles and the SO phase, predominantly in UP and transition to UP states, that relates to memory consolidation during sleep^39,72–74^. Based on this we explored the influence of cue time in relation to SO phases (and coupling with spindles) on decoding performance during NREM 2/3 sleep. In particular, we were interested in whether coupling might help further optimise model training to enhance its ability to learn semantic information during sleep, thereby improving decoding performance.

We categorised NREM 2/3 auditory-triggered epochs into eight non-overlapping SO phase bins (**Figure 5a**), which together accounted for approximately 40% of all NREM 2/3 samples (see **Methods** for details); the per-phase event counts are detailed in **Extended Data Table 5**. We used the trained SI-MD model to investigate inherent decodability of sleep data, and found for the “1^st^ Up” and “2^nd^ Up” categories, decoding results were 27.87±1.85% and 23.62±1.92% respectively, significantly and marginally significantly higher than for the “Down” category, which was 18.74±1.99% (“1^st^ Up”: *t*(11)=2.86, *P*=0.015; “2^nd^ Up”: *t*(11)=2.14, *P*=0.055, two-sided paired *t*-test). This accords with previous findings that decoding performance is better when the cue presentation is in the Up state of an SO, compared to the Down state^6^. Interestingly, the “Trans 1^st^ UP” category, with a decoding accuracy of 36.93±1.48%, outperformed the Up state categories (“1^st^ Up”: *t*(11)=6.35, *P*<0.001; “2^nd^ Up”: *t*(11)=6.26, *P*<0.001, two-sided paired *t*-test). The “Trans 1^st^ Up” category also showed significantly higher decoding accuracy compared to all other categories (“Pre SO”, “Trans Down”, “Down”, “Trans 2^nd^ Up”, “Post SO”: all *t*(11)>3.60, all *P*<0.01, two-sided paired *t*-test).

**Figure 5.**
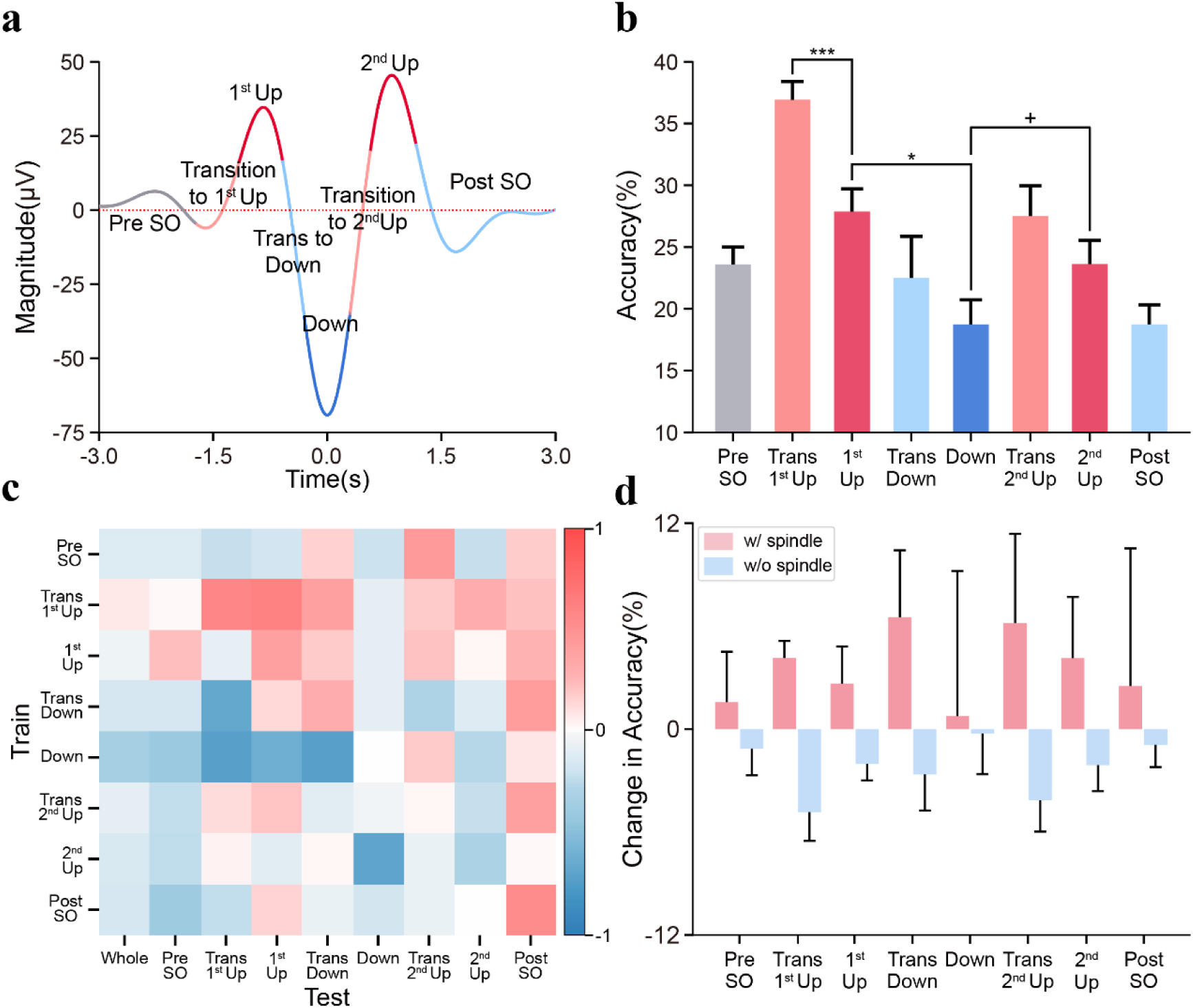
Sleep decoding performance as a function of slow oscillation phases and spindle coupling. (**a**) Slow oscillation (SO) phases. During NREM 2/3 sleep, we divided the phase of a SO event into eight parts (color-coded), based on the onset of audio stimuli. The time represented on the x-axis corresponds to a 6-second data length, with the trough of the SO event as the zero point. (**b**) Sleep decoding performance by SO phase. The decoding performance of different SO phases during NREM 2/3 sleep using the SI-MD model is shown with average performance and standard error across subjects. The decoding results for the “1^st^ Up” and “2^nd^ Up” categories were significantly and marginally significantly respectively higher than for the “Down” category, while the result for the “Trans 1^st^ Up” was significantly higher than those for the “1^st^ Up” and “2^nd^ Up” categories. The bar chart colors for all categories correspond to the colors in the diagram of panel (a). (**c**) Generalisation ability of trained sleep decoders across different SO stages. The same sleep decoder model was re-trained on an equivalent amount of data, separately for each SO stage during NREM 2/3 sleep, and compared to a baseline model, which was re-trained with an equivalent amount of training data randomly sampled from the SO phases, rather than within one of the specific categories. The change in decoding accuracy is color-coded: red indicates accuracy above the baseline model, and blue indicates accuracy below the baseline model. (**d**) Sleep decoding performance with versus without spindle coupling. Within each SO stage, we further compared the change in decoding performance with and without spindle coupling using the trained sleep decoder. This comparison was based on the presence or absence of a spindle event in the testing trials. The decoding performance for the “with spindle” group is shown in pink, while that for the “without spindle” group in blue. + *p*<0.10, * *p*<0.05, ** *p*<0.01, *** *p*<0.001.

Taking the differences in sample sizes in SO phases into consideration, we extracted equal amounts of data from each SO category in the training dataset to train the SI-MD model, then decoded the test dataset. The “Trans 1^st^ Up” was the sole category that showed results consistently above baseline across all eight categories except “Down” (**Figure 5c**), demonstrating a strong transfer generalisation ability.

We next explored whether SO coupling with sleep spindles influenced decoding ability. Temporal frequency analysis of the Fz channel waveforms (from -1.5 s to 3.0 s, based on cue onset) revealed significant coupling of the spindle (12-16 Hz) frequency band with SO phases, especially when the cue was in the “Trans 1st Up” stage of the SO (**Extended Data Figure 4**). Reapplying the trained SI-MD sleep model for decoding and using the overall decoding accuracy of the corresponding SO phase as baseline for each subcategory (**Figure 5d**; see also **Extended Data Figure 5** for the distribution of spindle coupling frequency and mean magnitude as a function of SO phases), we found that spindle coupling improved decoding accuracy across categories. Thus, the presence of spindle coupling significantly improved decoding accuracy for the “Trans 1^st^ Up” category, as well as for test subjects overall (“Trans 1^st^ Up”: *t*(11)=3.01, *P*=0.012; “Whole”: *t*(11)=4.48, *P*<0.001,two-sided paired *t*-test).

On this basis, we retrained the SI-MD model after removing training data corresponding to the ‘DOWN’ and ‘Post SO’ phases during sleep. To compensate for the reduced, and now ‘cleaned’ training set, we adjusted the model’s convolutional encoder accordingly (see details in **Methods** and **Extended Data Table 3**). We obtained a retrained SI-MD model that showed a further decoding improvement when confined to the ‘Trans 1^st^ Up’ part (40.02±1.99%) compared to the original SI-MD model (38.38±1.32%) with a significantly higher overall decoding accuracy: 23.45±0.62%(“Trans 1^st^ Up”: *t*(11)=1.93, *P*=0.079; “Whole”: *t*(11)=2.43, *P*=0.034 two-sided paired *t*-test).

### SI model generalisation for spontaneous and cue-driven memory reactivation in independent studies

To evaluate the robustness and practical utility of our SI model beyond the context of initial TMR-like paradigm, we assesed its performance across two independent experiments with distinct participant samples. Both experiments avoid using semantic informative cues during sleep, one even without any cueing during sleep to probe spontaneous memory reactivation, thereby addressing potential confounds introduced by cue delivery in the current SI dataset.

In Experiment 1 (**Exp. 1**; **Figure 6a**), we probed whether the SI model could detect spontaneous memory reactivations in the absence of auditory cues. We used a task design adapted from Schreiner, et al.^39^. Twenty-five healthy volunteers (mean age 23.80±0.61 years; 17 female) each completed two sessions – “apple” and “book” – separated by at least one week (see details in **Methods**). In each session, participants first learned forty image-verb associations (all images belong to the same conceptual category, “apple” or “book”, with the paired verbs being arbitrary, e.g., associating a “leafless red apple” image with the verb “run”), followed by a brief memory test. They then took a nap (79.5 ± 3.3 minutes; **Extended Data Table 6**) during which no sounds were played, and finally completed a post-sleep assessment.We treated the two concept categories as ground-truth labels, extracted NREM-2/3 epochs time-locked to slow-oscillation troughs, and applied the SI decoder to these segments (**Extended Data Figure 6**; see Methods).

**Figure 6.**
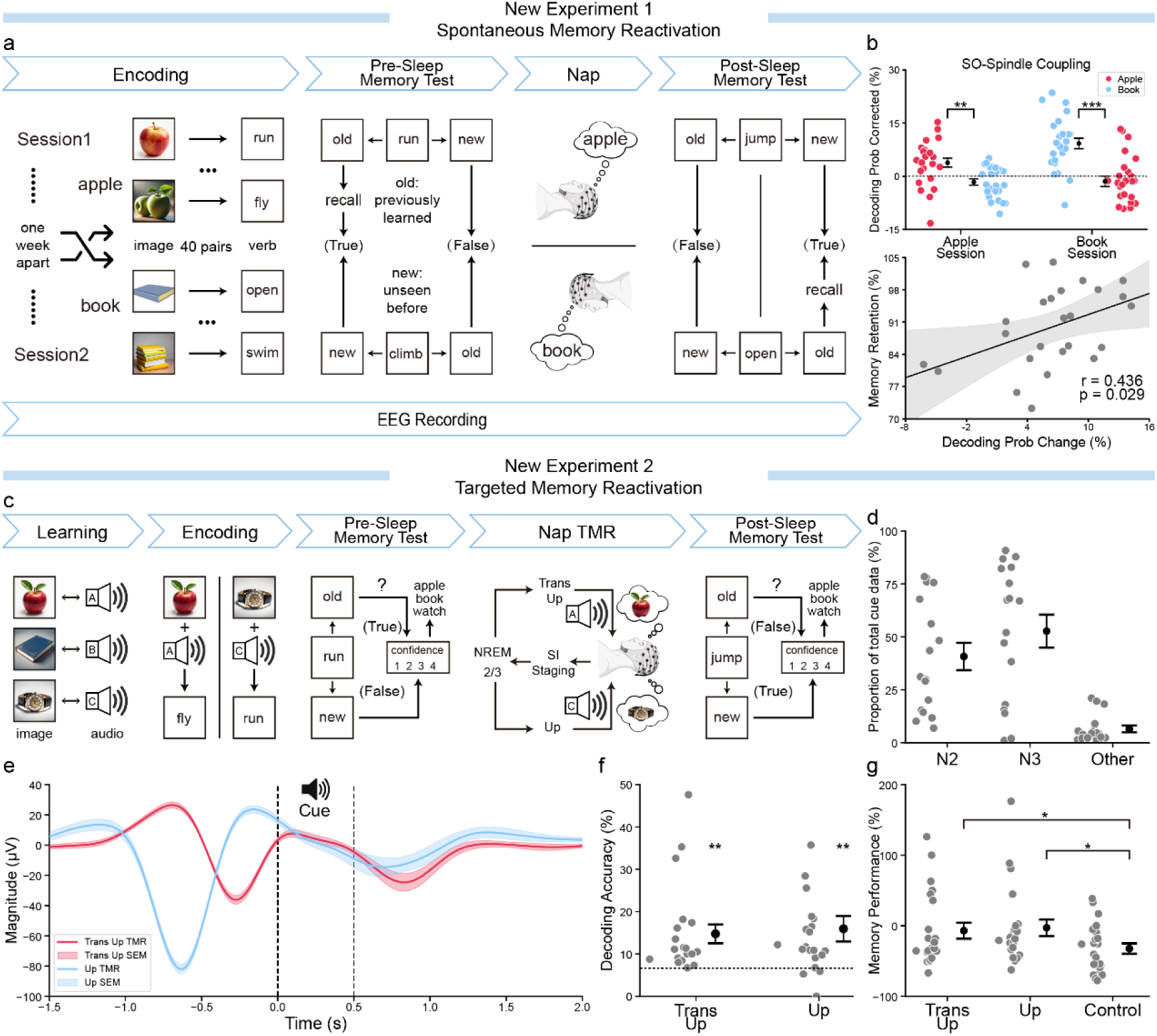
Generalisation of the SI model to independent datasets: Experiment 1 - spontaneous decoding without any TMR cues; Experiment 2 - decoding following TMR with non-semantic audio cues. **(a)** In New Experiment 1 (Exp. 1), two sessions were conducted more than one week apart, each centering on a single concept (“apple” and “book”) presented in random order. Participants paired 40 different images (each varying in features like number, colour, state, etc. but all belong to the same core concept, e.g., “apple”) with 40 distinct verbs (e.g., associating a “leafless red apple” image with the verb “run”). In the subsequent memory test, participants were presented with 20 randomly chosen verbs from the learned set and 10 entirely new verbs in random order; they judged each verb as “new” (unseen before) or “old” (previously learned). If a verb was judged “old”, they then typed a detailed description of the corresponding image, all without feedback. Immediately following this memory test, participants proceeded to the sleep phase, after which they completed a second memory test using the remaining 20 learned verbs and another 10 new verbs. Whole-brain 64-channel EEG recordings were collected throughout the experiment, during both sleep and awake tasks. More details can be found in **Methods**. **(b)** Experiment 1 results. SI decoding probabilities were computed for SO-spindle coupling epochs in NREM 2/3 and bias-corrected. In the “Apple” session, decoding favoured “apple”; in the “Book” session, it favoured “book”. Probabilities for the session-specific concept exceeded both the alternative concept and baseline, and correlated with memory retention (post/-pre accuracy). **(c)** New Experiment 2 (Exp. 2) is with TMR paradigm paired with semantic-free sounds and different images. Participants first learned three images representing three concepts (“apple”, “book”, “watch”, but different image as to the current SI dataset), each paired with a semantic-free sound. They then completed a 36-trial matching test with feedback, requiring at least 90% accuracy (“learning” phase. Note, this is different from the familiarisation phase in the main experiment, as in Exp. 2, subjects indeed need to learn the pairing between images and arbitrary sound which does not convey semantic information), before proceeding. In the subsequent encoding part, each image-sound pair (concept) was associated with 40 distinct verbs (total 120). A memory test then presented a total of 60 “old” verbs (20 per concept) plus 30 “new” verbs in random order, prompting participants to judge each verb as “new” or “old” and to rate their confidence. If a verb was judged “old”, participants were then required to identify which concept it belonged to before moving on, all without feedback. During the nap, real-time sleep staging was performed based on SI-Staging to detect NREM2/3 sleep, enabling the delivery of auditory stimuli for randomly selected concepts during either the “Up” phase or “Transition to Up” phase of the slow oscillations. After the nap, participants took a second memory test consisting of the remaining 60 learned verbs plus another 30 new verbs. Whole-brain 64-channel EEG recordings were collected throughout the experiment, during both sleep and awake tasks. More details can be found in **Methods**. **(d)** The proportions of cued data in N2, N3 and other stages of sleep, following offline, expert-based manual sleep staging. **(e)** ERPs at Fz for TMR are plotted in red for cues in “Transition to Up” phase and blue for cues in “Up” phase, shown as mean±SE across participants. Vertical dashed lines mark TMR cue onset and offset. (**f**) SI model decoding performance was significantly higher than chance level for TMR cues delivered during the “Transition to Up” and “Up” phase. The black dashed line denotes the 1/15 chance-level accuracy. **(g)** Post-sleep memory performance was evaluated for concepts cued during the “Transition to Up” phase, during the “Up” phase, and in an uncued control group. Performance in both the “Transition to Up” and “Up” cued conditions was significantly higher than in the control condition. The gray dots represent the results for each subject, while the black error bars indicate the mean (M) and standard error of the mean (SE) for the corresponding conditions. \**p*<0.05, ** *p*<0.01, *** *p*<0.001.

Across all sessions, decoding during SO-spindle coupling exceeded that of uncoupled segments (“Apple”: t(24) = 2.31, *P* = 0.029; “Book”: t(24) = 5.69, *P* < 0.001; **Extended Data Figure 7**). These decoding results were calibrated to offset any potential training-induced category bias (**Extended Data Table 7**). The model predicted probability of correctly classifying the session-specific concept during coupling also surpassed its probability for the alternative concept and the baseline reference (“Apple”: t(24) = 3.61, *P* = 0.001; “Book”: t(24) = 6.11, *P* < 0.001; baseline comparisons: all t(24) > 3.00, *P* < 0.01; **Figure 6b**). Moreover, the decoding probability during SO-spindle coupling predicted subsequent memory retention (post-/pre-sleep accuracy; *r* = 0.436, *P* = 0.029; **Figure 6b**). These findings show that the SI model, used off-the-shelf, can reveal spontaneous reactivation during sleep and that its outputs relate meaningfully to behavioural memory performance.

In Experiment 2 (**Exp. 2**; **Figure 6c**), we tested whether the SI model could decode reactivations triggered by semantically neutral cues. The task was adapted from that used in previous TMR research^6,40,54,55^.Twenty-two new participants (mean age 23.86±0.50 years; 15 female) first learned three images representing three concepts (“apple”, “book”, “watch”, but different image as to the current SI dataset), each paired with a distinct semantic-free sound. Subjects then learned pairings between arbitrary verbs and concepts, with each concept associated with 40 distinct verbs to ensure task difficulty and lexical diversity. There were one hundred and twenty verb–image pairs, followed by a brief memory test. They then underwent a nap (80.2±5.3 minutes; **Extended Data Table 6**) during which two of the three cues – balanced across participants – were delivered during NREM 2/3 sleep at either the “transition to up” or the “up” phase of the slow oscillation (SO), as detected from the Fz channel (**Figure 6d,e**; see also **Extended Data Figure 8**). A final post-sleep test assessed memory for both cued and uncued items.

Applying the SI model to cue-evoked sleep epochs, we observed decoding accuracies significantly above chance level (“Transition to Up”: *t*(21)=3.64, *P*=0.002; “Up”: *t*(21)=3.09, *P*=0.006, one-sample *t*-test; **Figure 6f**). Behaviourally, memory improvements from pre- to post-sleep were also greater for cued items relative to the uncued control condition (“Transition to Up”: *t*(21)=2.20, *P*=0.039; “Up”: *t*(21)=2.27, *P*=0.034, two-sided paired *t*-test; **Figure 6g**). These findings illustrate that our model can decode memory content even when the cues themselves carry no semantic information, and that cue-timing relative to the SO modulates both neural decoding and behavioural outcomes.

Together, results from both experiments confirm that the SI model generalises effectively to independent datasets, and can reliably detect both spontaneous and cue-induced memory reactivations. The correspondence between decoding accuracy, slow-oscillation dynamics, and memory retention highlights the model’s sensitivity to endogenous memory processes and its promise for real-world TMR and sleep-based memory research.

### Real-time sleep decoding system

Finally, we implemented a real-time sleep decoding system (**Figure 7**). EEG, EOG and EMG signals are acquired in 30 s epochs and pre-processed online before entering the SI-Staging model. Sleep-stage classifications are updated and displayed instantly in both the hypnogram and the staging panel. We measured the real-time latency of the SI-Staging model over 10,000 trials on various GPUs: 3060 Ti: 432.23±0.14 ms; 4060: 319.11±0.11 ms; 4070: 300.16±0.04 ms; 4080: 287.85±0.04 ms. Once a stable NREM 2/3 or RREM stage persisted for more than 30 s, the system automatically triggered the auditory-stimulus module, delivering sounds either randomly or according to a user-defined interval.

**Figure 7.**
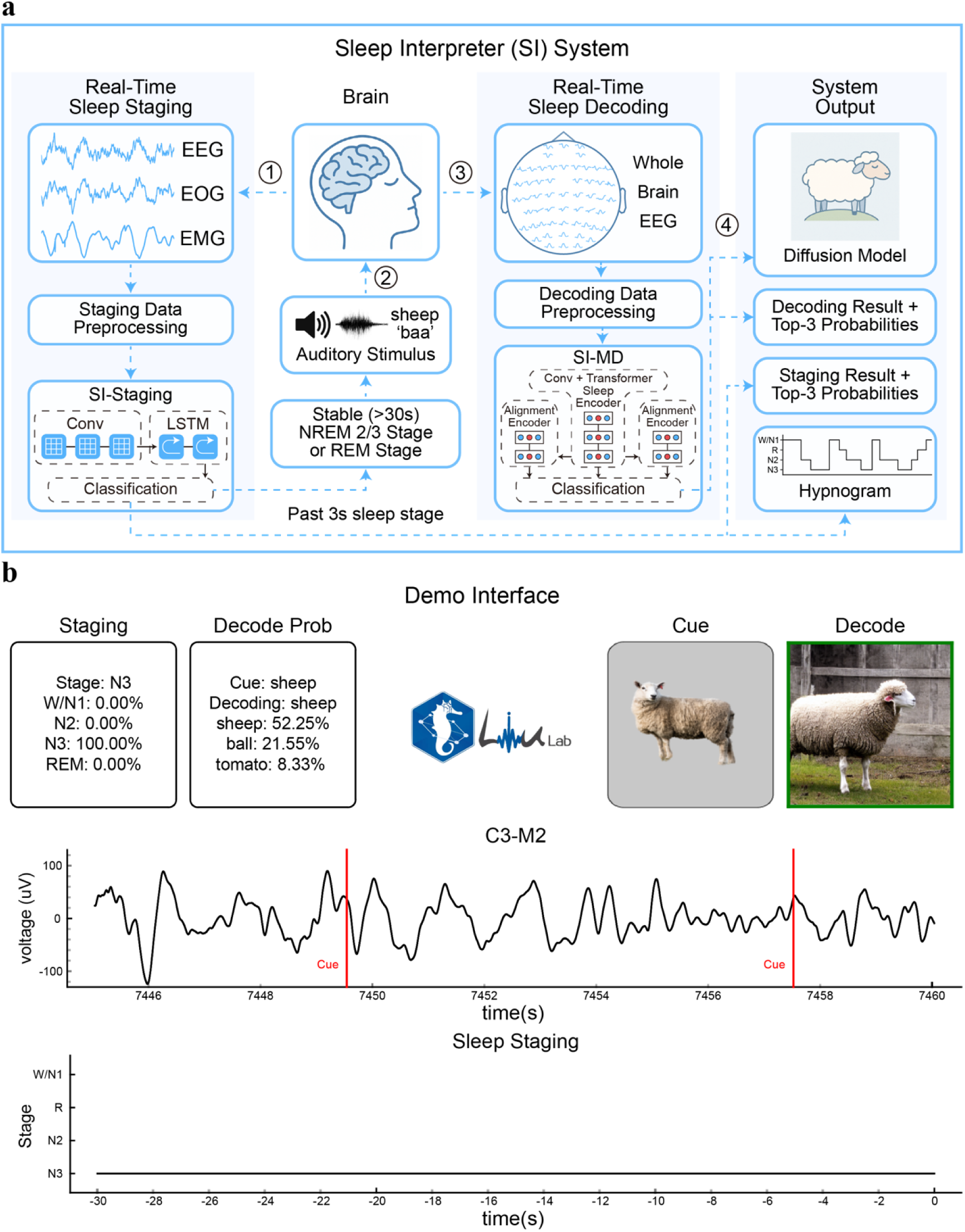
Data flow and interface of the real-time sleep decoding system. **(a)** Data flow. First, the neural signals, EEG from channels C3 and M2, plus EOG and EMG, are re-referenced, band-pass filtered, downsampled and normalised before entering the SI-Staging model. Then the sleep-stage probabilities are updated every 3 s and shown in both the hypnogram and staging panel. When a stable NREM 2/3 or REM stage persists for more than 30 s, the auditory-stimulus module delivers sounds (either at random or according to a user-defined schedule). Immediately afterwards,, online pre-processed whole-brain EEG is passed to the SI decoder; its semantic output is displayed in the decoding panel and also routed to a pre-trained diffusion model^75^ to generate a representative image. This image generation is for visualisation purpose only, e.g., to help an experimenter or potential user interpret or verify the decoded content in a real-time setting. **(b)** System interface. The top-left panels show the real-time sleep stage (left) and decoding result (right). The top-right panels display the last auditory-cue paired image (left) and the diffusion-model visualization driven by the decoder’s output (right). The middle panel presents the most recent 30 s of C3-referenced EEG (M2-reference; 0.1– 40 Hz), and the bottom panel shows the corresponding sleep stage for that epoch. See **Extended Data Video 1** for a live demonstration.

Subsequently, the whole-brain EEG is captured, pre-processed online, and streamed to the SI-MD decoder, whose output is shown in the decoding panel. Across 10, 000 trials, the SI model’s computation time was: 3060 Ti: 709.76±0.16 ms; 4060: 515.03±0.30 ms; 4070: 480.94±0.62 ms; 4080: 455.30±0.03 ms. Decoded class labels are also passed to a pre-trained diffusion model^75^ to generate a representative image, providing intuitive visual feedback (see Methods for details, see also **Extended Data Video 1** for a live demonstration).

By sharing this real-time sleep decoding platform, along with all accompanying code and data, our aim is to enable the wider community to adopt and build upon our system, thereby facilitating the development of novel sleep-based interventions for human learning and memory.

## Discussion

We introduced SI, a neural network that decodes memory replay from human sleep EEG, trained and validated on a dataset of 135 participants (∼400 hr awake; ∼1000 hr sleep) collected in a TMR-like paradigm. SI achieved up to 40.02 % Top-1 accuracy in a 15-way sleep classification on unseen individuals—far above the 1/15 chance level— demonstrating that semantic content can be reliably extracted from sleep EEG. We are releasing our full codebase, pretrained models, and both raw and preprocessed EEG data in the hope that this resource will be advance sleep research on memory and learning, as well as disorders linked to abnormal sleep.

We compared our SI model against five recent EEG-decoding architectures^68,70,76–78^ (**Extended Data Table 4**). Temporal-spatial “foundation” models^68,70,78^ excel when pretrained on very large EEG datasets, particularly for clinical tasks such as seizure detection; yet temporal models^76,77^, remain more effective in capturing the fine-grained neural dynamics of cognition. Our SI builds on such temporal-model paradigm— leveraging convolutional layers and Transformers for global context—and further augments it with a contrastive learning objective, yielding higher accuracy than prior CNN- or purely Transformer-based decoders.

At the heart of SI is a temporal contrastive loss. Contrastive learning is typically a self-supervised technique that trains models to pull semantically similar samples closer in latent space while pushing dissimilar ones apart^79–81^. This approach has been widely adopted in multi-modal architectures^43,65^. Here, we apply the same principle in a supervised context^46^ by aligning neural dynamics of the same memories across wake and sleep while separating different memories. This strategy enables SI to extract semantically relevant neural signals from noisy sleep recordings with the aid of wake-period examples, thereby mitigating issues of label noise.

We also subjected SI to two independent “off-the-shelf” validations. In New Experiment 1, participants took a no-cue nap, yet SI still decoded spontaneous reactivations whose probabilities tracked individual memory retention. In New Experiment 2, we delivered semantic-free sounds at specific slow-oscillation (SO) phases (“Transition to Up” and “Up” peaks), and SI’s decoded probabilities again correlated with post-nap memory performance. These replications provide confidence that our decoding is not an artefact of the original paradigm but generalises across tasks, stimuli, and subject samples.

Our results also reaffirm the critical role of SO-spindle coupling in memory processing, in line with previous studies^39,72–74,82^. For example, memory reactivation during NREM sleep—both spontaneous and cue-evoked—is stronger during the slow oscillation “Up” state than the “Down” state^6,55^. Consistent with this, we found that a specific phase at the transition into the Up state (“Trans 1st Up”) yielded the highest decoding performance of all, and was marked by the greatest incidence of SO-spindle coupling and the largest EEG amplitude. The subsequent “Trans 2nd Up” phase performed comparably to the canonical Up-state. By retraining SI with epochs containing SO-spindle coupling (i.e., the Up and Transition-to-Up phases), we further enhanced decoding performance, reaching an average Top-1 accuracy of 40.02%.

In New Experiment 1, sleep periods exhibiting SO-spindle coupling showed significantly higher decoding probabilities that closely tracked subsequent memory performance. Similarly, New Experiment 2 demonstrated that inducing reactivation with auditory cues was most effective when those cues were delivered during coupling. We anticipate that focusing future models on these high-yield coupling periods—for instance with self-supervised training—could further enrich the capture of semantic signals and improve sensitivity to latent memory content.

We would caution that SI currently decodes only a closed set of 15 items, and our TMR-like design—as is the case for classical TMR protocols^6,40,54,55^—cannot guarantee memory reactivation on every cue. Future expansions of our general approach might employ self-supervised techniques such as masked modelling^68,70^ or open-set contrastive learning^80,83,84^, alongside larger, more diverse stimulus libraries (e.g., ‘THINGS’ database^25,85,86^) to create a universal sleep decoder.

Finally, we would highlight that SI opens new avenues for closed-loop interventions: delivering cues precisely at optimal SO-spindle phases to enhance consolidation or to modulate maladaptive memories in clinical contexts. We also developed a preliminary SI-MD model for REM sleep, a state characterised by desynchronised, dream-associated activity^87–89^, which could, once expanded beyond the current categories, provide more objective insight into dream content, complementing traditional subjective reports^90,91^, although more work are needed to further validate its suitability.

In conclusion, we introduce a large, well-annotated EEG dataset for sleep memory decoding and, leveraging this resource, develop a real-time system that integrates automatic sleep staging with memory content decoding. We hope this approach and resource lay the groundwork for a truly universal sleep decoder, enabling more precise and effective sleep-based interventions, e.g., targeting the pathological memories that underlie nightmares.

## Methods

### Participants

We tested a total of 149 participants, with 14 participants excluded for a range of reasons: two due to excessive body movements and poor data quality, four because they did not fall asleep, four because of electrode issues (falling off or drifting), two due to snoring, one was interrupted by a wakefulness experiment, and one failed to complete the full wakefulness experiment. 135 participants (mean age 22.55±0.22; 72 female) remained after exclusions.

A separate cohort of 38 participants took part in the New Experiment 1. Of these, eight participants were excluded due to insufficient sleep duration (less than 20 minutes of NREM sleep), two participants were removed for failing to comply with experimental instructions, and three were excluded because of equipment malfunction during data acquisition. This resulted in a final sample of 25 participants (mean age 23.80±0.61; 17 female).

For New Experiment 2, 31 participants were recruited. However, two participants were excluded due to unstable sleep patterns and insufficient exposure to targeted memory reactivation (TMR) auditory cues (less than 25), four were excluded for poor behavioral performance, one additional participant was excluded due to excessively high task accuracy (95%), which prevented observation of potential TMR effects, and two more were excluded due to a failure in understanding the experimental task. Consequently, 22 participants (mean age 23.86±0.50; 15 female) remained for further analysis.

All participants were healthy nonsmokers with no color blindness or color processing deficiency, normal hearing, and no allergies to alcohol or scrubs. They maintained normal wake-sleep rhythms, had not permed or colored their hair in the last month, did not suffer from insomnia, problematic snoring, or nighttime sleep disturbances. All participants gave written consent prior to the experiment. The study was approved by the Chinese Institute for Brain Research, Beijing (CIBR) and conducted in accordance with the Declaration of Helsinki.

### Experiment Design

Our experiment paradigm comprised four stages: an initial familiarisation stage, a pre-sleep functional localiser stage, an overnight sleep targeted memory reactivation stage and a post-sleep functional localiser stage (**Figure 1a**). During the initial familiarisation phase, we used 15 concepts: *alarm*, *apple*, *ball*, *book*, *box*, *chair*, *kiwi*, *microphone*, *motorcycle*, *pepper*, *sheep*, *shoes*, *strawberry*, *tomato*, and *watch*. Each concept was paired with a semantically congruent picture and sound, yielding 15 distinct visual-audio stimuli pairs. Participants engaged in a self-paced exploration of these pairings (clicking on each image played its corresponding sound) and proceeded to the next stage only after affirming that they had fully mastered every correct pairing.

In the pre-sleep function localiser stage, subjects were instructed to perform two tasks: the image-audio task and the audio-image task. In the image-audio task, subjects were presented with 600 trials, with each trial containing one image stimulus and one audio stimulus, in image-audio order. We randomly selected one image stimulus and one audio stimulus from the set of image stimuli and the set of audio stimuli, respectively, where the image stimulus and the audio stimulus could be inconsistent (i.e., derived from different semantic classes). In total, we had 210 inconsistent trials and 390 consistent trials. The purpose of this design is solely to ensure that participants formed veridical and stable stimulus pair memories.

For the image-audio task, subjects were presented with the following study pipeline: a blank screen for 0.6 to 0.9 seconds (inter-trial interval), followed by a cross centered on the screen for 0.3 seconds, then an image stimulus for 0.8 seconds. After the image, a blank screen was shown for 0.3 seconds, followed by an audio stimulus for 0.5 seconds, and finally, another blank screen for 0.3 seconds. Then, subjects were tasked to indicate whether the presented image stimulus and audio stimulus were matched (i.e., coming from the same semantic class). Finally, we presented behavioral feedback for 0.2s. In total, the image-audio task took about 48 minutes.

The setup of the audio-image task was identical to that of the image-audio task, with the sole difference being that we presented image-audio pairs now in an audio-image order. As in the post-sleep function localiser stage, subjects were required to execute the same tasks in the pre-sleep function localiser stage.

In the overnight sleep target memory reactivation stage, we utilised a near real-time application of the YASA^58^ model which refers to use the past five minutes’ data in staging. We consider that a subject was in either NREM2/3 or REM stage only when the total probability of NREM2/3 or REM stage in the model output results exceeded 70%. As the staging result subject reached the N2/3 stage of NREM sleep, audio stimuli alone were presented every 4-6 seconds (the same as those used during wakefulness, lasting 0.5s), where these were randomly selected from the set of audio stimuli.

### New Experiment 1

In New Experiment 1 (**Exp. 1**), twenty-five healthy volunteers (mean age 23.80±0.61 years; 17 female) completed two sessions – labelled “apple” and “book” – separated by at least one week and counterbalanced in order. Each session comprised three stages: a pre-sleep memory phase, a nap, and a post-sleep memory test. This task design is adapted from Schreiner, et al.^39^.

A set of eighty images – forty apples and forty books – served as stimuli. The apple pictures varied in size (small vs. large), completeness (whole vs. partial), quantity (single vs. multiple), presence or absence of leaves, colour (red, yellow, green), and freshness (fresh vs. decayed); the book images varied in artistic style (cartoon vs. realistic), state (open vs. closed), quantity, colour and condition (new vs. old).

During the pre-sleep phase, participants first performed a five-minute psychomotor vigilance task (PVT) in which a central fixation cross was randomly replaced every 2-10 seconds by a counter that increased from 0 to 2 seconds in 20 ms steps; participants pressed the space bar as quickly as possible to stop the counter and received feedback on their reaction times (mean RT: 325.74±8.23 ms). They then completed a learning task to memorise correct image-caption pairings: each trial began with a fixation cross (1.5±0.1 s), followed by a image with a descriptive caption shown for 2.5 s; forty image-caption pairs with accurate descriptions were retained for later tasks, while ten incorrect pairings were excluded. Next, in the encoding task, participants learned forty arbitrary image-verb associations: a fixation cross (1.5±0.1 s) preceded a verb (1 s) and then the corresponding image (4 s); participants were asked to imagined a scenario linking the verb to the image and, within 10 s of image offset, indicated whether the imagined scenario was realistic or bizarre. This block was repeated four times in varying orders to ensure robust encoding. Finally, a pre-sleep memory test presented twenty “old” (learned) verbs and ten novel “new” verbs: each trial began with a fixation cross (1.5±0.1 s) followed by a verb (3 s), after which participants made an “old” or “new” judgement. A “new” response ended the trial; an “old” response prompted a typed description of the associated image (or “do not know”), with accuracy scored against the original captions.

Participants then took a nap lasting 79.5±3.3 minutes on average, during which 64-channel EEG was continuously monitored with no sounds played. Twenty minutes after awakening, they repeated the PVT (mean RT: 326.31±6.11 ms; no significant difference from the pre-sleep session, *P*=0.93, two-sided paired *t*-test) and completed a second memory test on the remaining twenty learned items, following the same procedure as in the pre-sleep test.

### New Experiment 2

In New Experiment 2 (**Exp. 2**), twenty-two new participants (mean age 23.86±0.50 years; 15 female) completed a three-stage TMR nap study using semantically neutral cues. The task was adapted from previous TMR research^6,40,54,55^.

The pre-sleep memory phase began with learning three concept images (“apple,” “book” and “watch”), each paired with a unique semantic-neutral sound stimulus. Participants then encoded 120 verb-image pairs (forty verbs per concept) by forming vivid mental scenarios while their associated auditory cue (500 ms duration) played. Then, in the pre-sleep memory test, participants judged sixty encoded verbs (20 verbs per concept, for 3 concepts) and thirty novel verbs: after a fixation cross (1-1.5 s) and a verb presentation (3 s), they indicated “old” or “new,” rated confidence on a four-point scale, and for “old” items selected the corresponding concept or “forgotten” (within 10 s).

During the nap (80.2±5.3 minutes), our real-time SI-Staging model classified sleep stages using Fz for capturing candidate SO events, C3 for standard staging, and M2 as reference. Upon detecting stable NREM 2/3 (probability ≥ 0.7 for ≥ 30 s), a 0.5-2 Hz bandpass filter and Hilbert transform computed instantaneous phase and slope at Fz, ensuring auditory stimulation at either the 270° “transition to up” zero crossing or the 0° “up” peak of the slow oscillation (SO). Two of the three cues – balanced across participants – were delivered at these phase points (average 78.64±10.26 cues per condition), set to 42 dB.

Twenty minutes after waking, participants completed a final memory test on the remaining sixty encoded verbs and another thirty new verbs, using the same procedure as the pre-sleep test. This design allowed us to assess the SI model’s ability to decode both spontaneous and cue-driven memory reactivation across independent samples and recording conditions.

### EEG Data Recording

In the main study, continuous electroencephalography (EEG) data were acquired using a BrainVision 64-channel elastic cap, arranged according to the international 10-20 system. The online reference was set to FCz, and electrode impedances were kept below 5 kΩ. A bipolar EOG montage was established by positioning one electrode below the left eye and another above the right eye; three additional electrodes were placed on the chin and bilaterally beneath the chin to record electromyography (EMG). During the functional localisation task, signals were sampled at 500 Hz with no online filtering, whereas during overnight sleep, EEG were filtered online at 0.1-35 Hz and both EOG and EMG at 10-100 Hz.

### New Experiments 1 and 2

In both experiments, EEG was recorded with an ANT 64-channel system that likewise adhered to the 10-20 standard. Impedances were maintained below 10 kΩ, EEG signals were referenced online to CPz, and data were sampled at 500 Hz. Placement of EMG, ECG, and EOG electrodes mirrored that of the main study.

### Sleep Staging Data

We followed the standard pipeline of YASA^58^. Accordingly, we provided a central electrode C3 referenced to the mastoid M2 (i.e., C3-M2) as the EEG channel, the right EOG channel REOG referenced to the mastoid M2 (i.e., REOG-M2) as the EOG channel, and the right EMG channel REMG referenced to the reference EMG channel EMGREF (i.e., REMG-EMGREF) as the EMG channel. For each sleep staging dataset, we have expert-labeled manual staging results, which divide the data into 30-second stages. Based on these manual staging results, we first calculated the Total Sleep Time (TST), Sleep Efficiency (SE) and the proportion of each stage (i.e., Wake, N1, N2, N3 and REM stages) to confirm the sleep structure, then segmented the data into 30-second intervals and augmented it with a sliding window of 3-second length. For data segments containing multiple labels, we assigned the label that accounted for the majority of the segment as the corresponding segment label. Finally, we used the ‘scale’ function of the ‘preprocessing’ module in the python package ‘sklearn’ for normalisation.

### Sleep Decoding Data

We used the python package MNE^92^ to epoch the sleep task-related EEG signals from -0.2 s to 1.8 s according to stimulus onset, dropping noisy epochs automatically, via “mne.epoch” function. Additionally, for sleep EEG signals, only data in the manually labeled NREM2/3 and REM stages were taken out. It should be noted that three subjects had no usable REM data due to the absence of stimuli within their short REM sleep. Each channel of the data segments was then re-referenced by subtracting out the mean signal over all EEG channels (average re-referencing). For data segments in REM stage of sleep, these were filtered within the frequency range of 0.1-40Hz using ‘filtfilt’ function in python package ‘signal’ (bandpass, order=4), while line noise at 50Hz (and its harmonics) of data segments in REM stage of sleep were removed using ‘iirnotch’ function in python package ‘signal’. Finally, the preprocessed data segments of NREM2/3 and REM stages were all resampled to 100Hz using the ‘decimate’ function in python package ‘signal’.

The next preprocessing for awake EEG signals was performed using the python package MNE^92^. The awake EEG signals were filtered within the frequency range of 0.1-40Hz (bandpass, order=4). Then, line noise at 50Hz and its harmonics were removed. Each channel was re-referenced by subtracting out the mean signal over all EEG channels (average re-referencing). We then applied Independent Component Analysis (ICA) that removed artifact components automatically^93^. The preprocessed EEG signals were further resampled to 100Hz. Finally, we epoched the EEG signals from -0.2 s to 0.8 s according to the onset of the stimuli cue (e.g. image, audio), and dropped noisy epochs automatically.

Next, in python we applied the ‘robust_scale’ function from the ‘preprocessing’ module of the ‘sklearn’ package to normalise all the preprocessed awake and sleep data segments, clamping the normalised values greater than 10 and less than -10. For the awake training data, we randomly removed half of the data from the same domain with the same labels for each individual, averaged the remaining data to obtain a new awake data segment, and generated a new awake dataset of the same size as the corresponding sleep dataset for model input.

### Real-time Sleep Staging

Following our sleep staging data preprocessing, we formulated the EEG signals as X ∈ ℝ^*C*×*T*^, where *C* is the number of EEG channels and *T* is the total timestamps. The associated label was denoted as *y* ∈ *Y*, where *Y* represents the set of pre-determined stages. In summary, for each subject, the total dataset was *D*_*s*_, *s* ∈ *S*, where *S* is the set of subjects and each sub-dataset comprised paired EEG-concept data (⟨*X*, *y*⟩). Therefore, the goal was to decode the corresponding stage label *y* from the preprocessed EEG signals.

For optimal staging efficacy, the model harnesses data from three channels (EEG, EOG, EMG) as inputs. For each channel, the input data has a sampling rate of 500 Hz and a total length of 30 seconds, which corresponds to X ∈ ℝ^3×15000^. To meet the requirements of real-time sleep staging, we set the input data with a sliding window applied every 3 seconds for subsequent staging. For the staging results of the real-time sleep model, we considered the initial 30-second staging results as the onset of the initial sleep state. Subsequently, based on this onset, we assigned the staging results after each sliding window to the corresponding sleep state within the 3-second time period. We trained the model in a supervised manner, predicting four sleep stages: W/N1, N2, N3, and REM. The corresponding labeling values for these stages are 0, 1, 2, and 3, respectively.

Our SI-Staging model consists of two main parts: the CNN module and the LSTM module. Through the tandem processing of the CNN and LSTM modules, followed by a final classification layer, we obtained the predicted label *y*′. The loss term of the model encompasses cross-entropy error and mean square error calculated between the ground truth label *y* and predicted label *y*′:

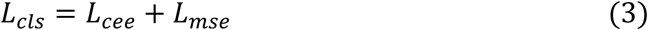

When calculating the cross-entropy error, the ground truth label and predicted label values are converted into one-hot vectors. For the mean square error calculation, they are treated as simple numbers.

The staging model was trained up to 200 epochs by default using Adam with a learning rate of 3 · 10^−4^ and a batch size of 256 on one NVIDIA V100 GPU with 32GB of memory.

### Within-domain Decoding and Cross-domain Generalisation Analysis

After preprocessing, each EEG segment was represented as X ∈ ℝ^*T*×*C*^, where *C* is the number of EEG channels and *T* is the number of time points.

#### Within-domain decoding

For each state (wake or sleep) we trained a Lasso-regularised GLM with 5-fold cross-validation on individual participant data, using one binary model per class across the 15 semantic labels. Classification probabilities were obtained with a softmax and converted to predicted labels via argmax; accuracy was then averaged across all test samples to yield the within-domain score.

#### Cross-domain generalisation

Each 2 s sleep epoch was split into 13 overlapping 0.8 s windows (0–0.8 s, 0.1–0.9 s,…, 1.2–2.0 s) to match the length of wake epochs. For every window we trained Lasso-GLMs (one per class) on all channels (awake task evoked, or sleep), and applied them to the corresponding 0.8 s segment in the other domain. For the decoding outcome, we took the mean accuracy over all computed time windows to obtain the cross-domain score.

#### Permutation testing

To evaluate significance, we reran the entire pipeline 1 000 times with randomly shuffled class labels, generating null distributions for both within- and cross-domain accuracies. The 95th percentile of the permuted distribution served as the significance threshold, minimising the risk of spurious effects from any single permutation.

### Real-time Sleep Decoding Model

We formulated the awake or sleep EEG signals after sleep decoding data preprocessing as X ∈ ℝ^*T*×*C*^, where *C* is the number of EEG channels and *T* is the total timestamps. The associated semantic label was denoted as *y* ∈ *Y*, where *Y* represents the set of 15 pre-determined semantic concepts (each paired with both an audio stimulus and an image stimulus), stored in the form of one-hot encoding. In summary, for each subject, the total dataset contained three sub-datasets, i.e., 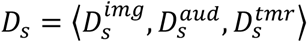, *s* ∈ *S*, where *S* is the set of subjects and each sub-dataset comprises paired EEG-concept data (⟨*X*, *y*⟩). Therefore, the goal was to decode the corresponding semantic label *y* from TMR-related raw EEG signals *X*^*tmr*^, or multidomain EEG signals _⟨ *X*_*img*_, *X*_*aud*_, *X*_*tmr*_⟩._

Our SI-Multi Domain (SI-MD) model contains two parts: (1) Convolutional Encoder, which includes one-dimensional convolutional layers and a linear projection module; (2) Transformer Encoder (see full configuration details in **Extended Data Table 3)**.

(1) Convolutional Encoder. As each EEG sample *X* has multiple channels, it is vital to fuse multiple channels to extract meaningful tokens before token-wise interaction by self-attention. Our convolutional encoder, which comprises a convolution module with residual convolution blocks and a linear projection module, encodes raw EEG signals into token embeddings. Each residual convolution block contains two 1-D convolution layers and a batch normalisation layer^94^ and the linear module contains two linear projection layers. We denote the output token embeddings from the convolutional encoder as

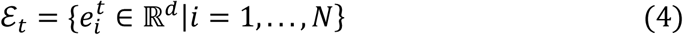

where *d* is the dimension of the embeddings, and *N* is the number of token embeddings.

(2) Transformer Encoder. Before feeding the embedding sequence into the transformer encoder, we add the parameter-free position embeddings introduced in previous work^64^ to enable the model to be aware of the relative positions among embeddings. Then, the sequence of embeddings will be fed into the transformer encoder to get the final encoded ℰ = {*e*_*i*_ ∈ ℝ^*d*^|*i* = 1,…, *N*}. For the downstream 15-way classification task, we flatten the output embeddings followed by a classification layer.

Since the neural patterns differ greatly among different tasks, we utilise three different convolutional encoders to extract meaningful tokens from raw EEG signals ⟨ *X*^*img*^, *X*^*aud*^, *X*^*tmr*^⟩. After tokenisation, the embeddings are fed into the transformer encoder for sequence modeling. Then, the output embeddings ℰ are used to predict the semantic label ŷ and calculate the classification loss.

To better utilise the template from the awake dataset, we introduce two special model designs: (1) a contrastive objective to align awake and sleep neural latent sequence position-wise, and (2) a alignment encoder to further improve alignment. The overall architecture of model is shown in **Figure 4a**.

#### Contrastive Objective

We calculate contrastive loss and align the neural representation sequences sharing the same category, originating from different tasks. Our contrastive loss consists of three parts, each aligning representations from two different tasks.

For each batch ℬ, we randomly draw|ℬ|sample pairs ⟨*X*^*tmr*^, *y*⟩ from the whole TMR-related dataset 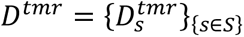. Then, for each sample pair, we further draw one image-evoked sample *X*^*img*^and one audio-evoked sample *X*^*aud*^ according to the semantic label *y* and the subject id *s*. We have 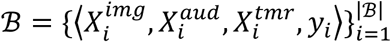, where different items could come from different subjects. After mapping the raw EEG signals *X* into encoded embedding sequence ℰ, we utilise the position-wise projection to transform embeddings and flatten them to form the latent embedding *z*. Then, we calculate the contrastive loss of three combinations based on the Equation-1.

#### Alignment Encoder

In our experiments, directly aligning the neural representation sequence of TMR-related EEG signals *X*^*tmr*^ with that of both *X*^*img*^ and *X*^*aud*^ often led to a slight degradation, compared to aligning with either of them individually. The initial design required the model to simultaneously find the optimal alignment among them, which posed a challenging task. To further smooth the aligning process, we feed the output embeddings ℰ^*tmr*^ from *X*^*tmr*^ into the transformer encoders of awake signals, yielding ℰ^*tmr*2*img*^ and ℰ^*tmr*2*aud*^ respectively. Therefore, we utilise the transformed embeddings to calculate contrastive loss, i.e., ℰ^*img*^ vs. ℰ^*tmr*2*img*^ and _ℰ_*aud* _vs. ℰ_*tmr*2*aud*.

Besides, all of these embeddings ⟨ℰ^*tmr*^, ℰ^*tmr*2*img*^, ℰ^*tmr*2*aud*^⟩ are utilised to predict the semantic labels, and averaged to get the final prediction *y*, which resembles the model ensemble method.

The total loss for training model consists of two parts: (1) classification loss, and (2) contrastive loss. For each data item (*X*^*img*^, *X*^*aud*^, *X*^*tmr*^, *y*), all of these transformed embeddings ⟨ℰ^*tmr*^, ℰ^*img*^, ℰ^*aud*^, ℰ^*tmr*2*img*^, ℰ^*tmr*2*aud*^⟩ are utilised to calculate the classification loss:

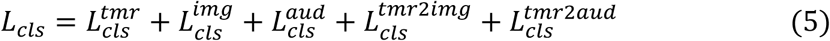

And the contrastive loss is calculated as demonstrated before in Equation-2: Therefore, the total training objective of model is:

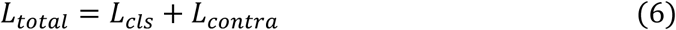

The decoding model was trained up to 100 epochs by default using AdamW with a learning rate of 8 · 10^−5^, a weight decay of 2 · 10^−5^ and a batch size of 256 on four NVIDIA V100 GPU with 32GB of memory.

### Decoding Model Comparison and Ablation Experiment

The five models employed for comparison are as followed: the Logistic GLM model, the LSTM-based model, the CNN-based model, the Transformer-based model, and our Sleep Interpreter – Single Domain (SI-SD) model. The Logistic GLM follows the standard setup in neuroscience: we train and evaluate a time-specific classifier for each time point, then take the maximum accuracy among these classifiers. The LSTM-based model comprises two basic LSTM layers, while the CNN-based model comprises two basic convolutional layers. The Transformer-based model consists of transformers with a depth of 8 and an attention head count of 6. For the SI-SD model, it includes the same architecture as the SI-MD model but excludes the audio encoder, the image encoder, the convolutional encoders of awake data domains and neural contrastive learning. All models used for the comparison were ensembled using five same structure models^95,96^.

### Pretrain-finetune Pipeline

Our method consists primarily of two stages: the pretraining stage and the finetuning stage. During the pretraining stage, we leave one subject out, e.g. subject *i*. Then, we use the datasets from the rest subjects to format the training dataset 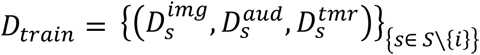. Following the previous training procedure, we get a pretrained model for that subject.

The fine-tuning process comes after the pretraining stage, in which we will finetune the pretrained model using data from the test subjects. For each dataset of test subject *i*, i.e., (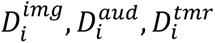), we split the dataset into training, validation, testing splits with a size roughly proportional to 80%, 10%, and 10%. We train each model with the training split for 200 epochs, by default using Adam with a learning rate of 2 · 10^−5^ and a batch size of 64 on one NVIDIA V100 GPU with 32GB of memory. And then evaluate its performance on validation and testing splits. We take the test accuracy according to the maximum validation accuracy as its performance and compute their average as their performance.

### Slow Oscillation Phases

Based on the criteria proposed by existing studies^97^, we detected the “Slow Oscillation” (SO) events.

First, raw EEG data was filtered between 0.16–1.25 Hz using ‘sosfiltfilt’ function of python package ‘signal’ (bandpass, order=4). Second, all zero-crossings were determined in the filtered signal, and event duration was determined for SO candidates (Down states followed by Up states) as time between two successive positive-to-negative zero-crossings for Fz channel. Events that met the SO duration criteria (minimum of 0.8 and maximum of 2 s, 0.5–1.25 Hz) entered the next step. Third, event amplitudes were determined for the remaining SO candidates (trough-to-peak amplitude between two positive-to-negative zero crossing for Fz channel). Events that also met the SO amplitude criteria (≥75% percentile of subject’s overall amplitudes, that is, the 25% of events with the largest amplitudes) were considered SO events.

Then, we segmented the same raw EEG signals into 6-second samples from -2.0 s to 4.0 s according to the onset of audio stimulus using python package MNE. We categorised each cue-triggered sleep epoch by SO phase, using the first SO event detected in that epoch. Phases were defined as follows:

1. Pre SO: This phase refers to the onset of the audio stimuli occurring before the 1 s interval immediately preceding the start of the first up-state of the SO. To be more precise, this phase should be called the “Pre Trans 1^st^ Up phase”.
2. Trans 1^st^ Up: This phase refers to the onset of the audio stimuli occurring during the transition to the “1^st^ Up” phase of the same SO event. Specifically, the transition corresponds to the onset of audio stimuli located within 1s prior to the start point of the detected “1^st^ Up” phase.
3. 1^st^ Up: In the time range of one second before the start point of the detected SO event, there could exist a waveform similar to the conventional “Up” state^6,97^. And there is another zero-crossing point in this time range, whereby we can determine a wave peak and take the portion above 50% of the peak value as the “1st Up” phase. If the required zero-crossing cannot be detected in the corresponding one second time period, we choose to use the maximum of the wave peaks in this period as the reference value for division.
4. Trans Down: This phase refers to the onset of the audio stimuli occurs during the transition to the “Down” state of the detected SO event. The transition corresponds to the stage between the end of the “1^st^ Up” phase and the beginning of the “Down” phase.
5. Down: The “Down” phase aligns with the “Down” state in previous studies^6,97^. However, we further refine this by strictly defining the “Down” phase as the portion below 50% of the trough value, and the onset of the audio stimuli occurs during this portion corresponds to the “Down” phase.
6. Trans 2^nd^ Up: This phase refers to the onset of the audio stimuli occurs during the transition to the “2^nd^ Up” phase of the same SO event. This corresponds to the stage between the end of the “Down” phase and the beginning of the “2^nd^ Up” phase.
7. 2^nd^ Up: The “2^nd^ Up” phase aligns with the “Up” state in previous studies^6,97^, but we define it more strictly. Only the onset of the audio stimuli occurs during portions of the former Up state waveform that exceed 50% of the peak value corresponds to the “2^nd^ Up” phase.
8. Post SO: This phase refers to the onset of the audio stimuli occurring after the SO event. Instead of using the traditional zero point, we define the end of the SO as the end of the 2^nd^ Up phase of the same SO event.

### SO-Spindle Coupling

The detection method for spindles and spindles coupling was as follows. First, raw data was filtered between 12–16 Hz using ‘sosfiltfilt’ function of python package ‘signal’ (bandpass, order=4). The mean amplitude was then calculated for the filtered signal using a moving average of 200 ms, with the spindle amplitude criterion defined as the 75th percentile of the subject’s overall mean amplitude values. Finally, whenever the signal exceeded this threshold for more than 0.5s but less than 3s (duration criteria), a spindle event was deemed to have been detected.

For the detected spindles, we used the trough point of the cue-evoked wave in the segmented sample as the starting point for SO-spindle coupling judgment. To simplify the analysis, temporal-frequency analysis showed that the low-frequency signal activation representing the evoked wave appeared 0.8s after the sound stimulus onset and the crossing after trough appeared around 1.2s after the sound stimulus onset. Since our decoding analysis only included data up to 1.8 s after the sound stimulus onset, spindle detection extended to 1.8 s after the sound stimulus onset to ensure that the analysis results match the existing decoding results. Thus, the detection time range for spindle-coupling was defined as 1.2-1.8 s after the sound stimulus onset.

### Temporal-Frequency Analysis

Temporal-frequency plots were calculated per event epoch (−2.0 s to 4.0 s according to the onset of audio stimulus) on Fz channel of raw data, using the ‘STFT’ function of the ‘signal’ package in python. The frequencies ranged from 0.1 Hz to 50 Hz in 13 steps, utilizing a sliding Blackman window with a variable, frequency-dependent length that always comprised 12 cycles. Time-locked temporal-frequency plots of all epochs were then normalised as percent change from the pre-event baseline (−2.0 to −1.5 s according to sound stimulus onset) and averaged per participant. The entire temporal-frequency analysis was conducted on Fz channel of the same sleep EEG data used for SO detection.

### Frequency Band Analysis

We filtered the raw EEG data using the following ranges: 0.5-4 Hz, 4-8 Hz, 8-12 Hz, 12-30 Hz, and 30-80 Hz, for delta, theta, alpha, beta, and gamma bands, respectively using ‘filtfilt’ function of python package ‘signal’ (bandpass, order=4). For the filtered data, we pretrained the SI-MD model separately with the training data filtered by each of the five frequency bands and used the pre-trained models to test on a total of six types of test data (one unfiltered and five filtered).

### Magnitude Analysis

We selected the raw sleep EEG data based on the Fz channel and calculated the mean magnitude within the range of -0.2 to 1.8 s relative to the sound stimulus onset. The mean magnitude was extracted using the Hilbert transform and convolved with a time window of 0.2s. The calculation range was adjusted to -0.15 to 1.75 s to account for the 0.2s sliding window and to avoid interference from out-of-bounds data.

### Model Retrain

We removed the data in “Post SO” phase and data in “Down” phase without SO-spindle coupling. Subsequently, we adjusted the depth of the convolutional encoder in SI-MD model structure. Considering the reduced size of the training dataset, we reduced the depth of convolution module from three residual convolution blocks to one block while maintaining the linear module of the convolutional encoder in SI-MD model (see detailed configuration in **Extended Data Table 3)**. Finally, we retrained the adjusted SI-MD model using the cleaned data. The model was trained up to 100 epochs by default using AdamW with a learning rate of 8 · 10^−5^, a weight decay of 2 · 10^−5^and a batch size of 256 on four NVIDIA V100 GPU with 32GB of memory.

### Spontaneous Memory Decoding

In New Experiment 1, we decoded spontaneous memory reactivation by time-locking the EEG to each slow-oscillation (SO) trough. For every trough we extracted a 4 s segment (−2 s to +2 s) and applied the SI-MD decoder with a 2 s analysis window, advanced in 0.01 s steps across the segment. To prevent bias, segments were randomly divided into validation and test sets before decoding. The 2 s window that yielded the highest decoding probability in the validation set was then applied to the held-out test set and to segments containing SO-spindle coupling.

To compensate for potential decoding biases introduced during model training, we established a calibration baseline using sleep data from the main dataset. We selected spontaneous SO epochs in NREM 2/3, excluding periods from 4 s before SO onset to the end if auditory stimuli occurred, and further retained epochs that met the SO-spindle coupling criterion. These data were segmented and decoded with SI-MD in the same way (**Extended Data Table 7**). The mean decoding probability across all trials served as the baseline for correcting decoding outcomes for both SO-only and SO-spindle coupling epochs in Experiment 1.

### Latent Diffusion Model

To visualise decoded sleep representations, we used an off-the-shelf text-to-image model: Stable Diffusion v1-5 (model card ID: runwayml/stable-diffusion-v1-5^75^). The SI decoder outputs a semantic concept label for each audio cue, which we embed into a fixed prompt designed to evoke a dream-like style (e.g. “softly blurred, dreamscape, impressionistic brushwork, cinematic bokeh”). No additional training or fine-tuning of the diffusion backbone was performed. For each epoch, a single image was generated using 50 denoising steps and a guidance scale of 7.5. These images provide an illustrative, abstract visualisation of sleep content rather than high-fidelity photographic reconstruction, and thus no explicit resolution constraints were imposed.

## Acknowledgment

Conceptualisation, Y.L., Z.C., H.Z., J.Z., L.Z., R.D., and T.B.; Investigation, Z.C., H.Z., J.Z., L.Z., P.L., H.W., Y.L., and T.B.; Writing – Original Draft, Z.C., Y.L.; Writing – Review & Editing, Y.L., Z.C., R.D., and T.B. This study is supported by the National Science and Technology Innovation 2030 Major Program (2022ZD0205500), the National Natural Science Foundation of China (32271093), the Beijing Natural Science Foundation (Z230010, L222033), and the Fundamental Research Funds for the Central Universities. R.D. and Z.C. are supported by UK DRI Key Questions Funding Programme (DRI-KQ2024-S1-1).

## Data availability

The benchmark sleep-TMR dataset will be available on https://osf.io/x7453/ upon publication.

## Code availability

The code for real-time sleep decoding system will be available on https://gitlab.com/liu_lab/sleep-interpreter upon publication.

## Extended Data Information

**Extended Data Figure 1.**
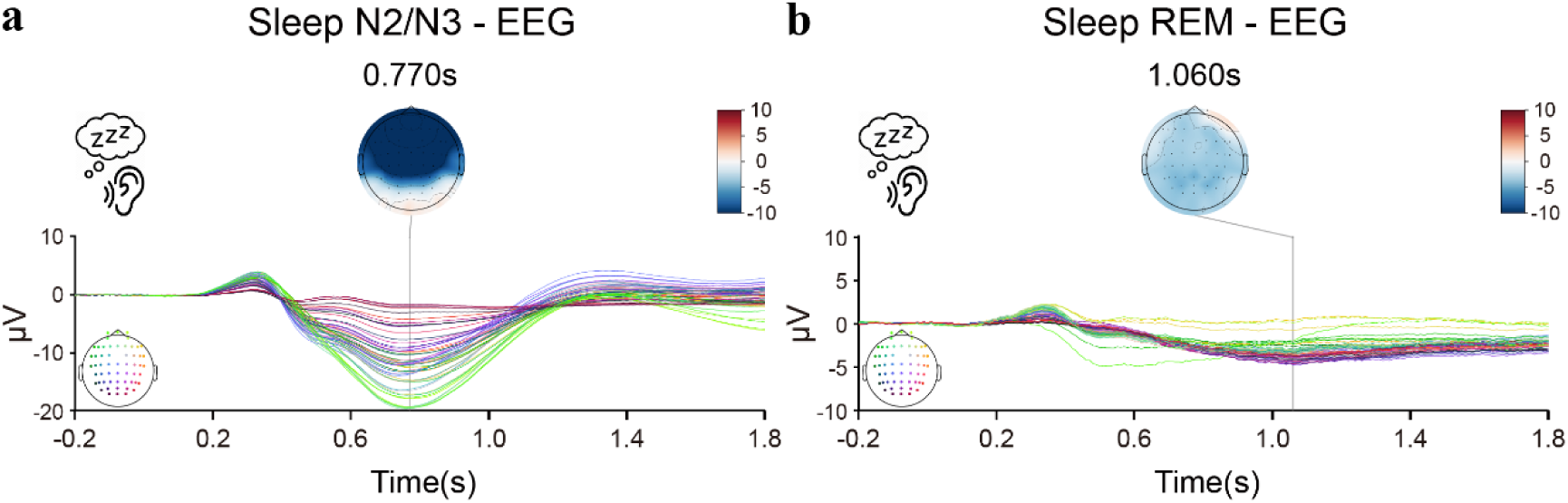
The averaged event-related potentials (ERPs) and scalp topographies evoked by audio cues during NREM 2/3 and REM sleep, derived from real-time preprocessed EEG (0.1-40 Hz bandpass filter). (**a**) ERPs and corresponding topographic maps evoked during NREM 2/3 sleep. These traces and maps illustrate the neural response in NREM, emphasising the typical slow-wave morphology. (**b**) ERPs and corresponding topographic maps evoked during REM sleep. These recordings, shown without offline artefact correction, highlight residual eye-movement effects characteristic of REM, thereby providing a direct visual contrast with the cleaner NREM responses. Time zero on the horizontal axis marks audio cue onset; ERPs were computed from -0.2 s to 1.8 s relative to onset. Topographic maps are displayed at the ERP peak latencies, using a common amplitude scale (−10 to +10 µV) to facilitate direct comparison across sleep stages.

**Extended Data Figure 2.**
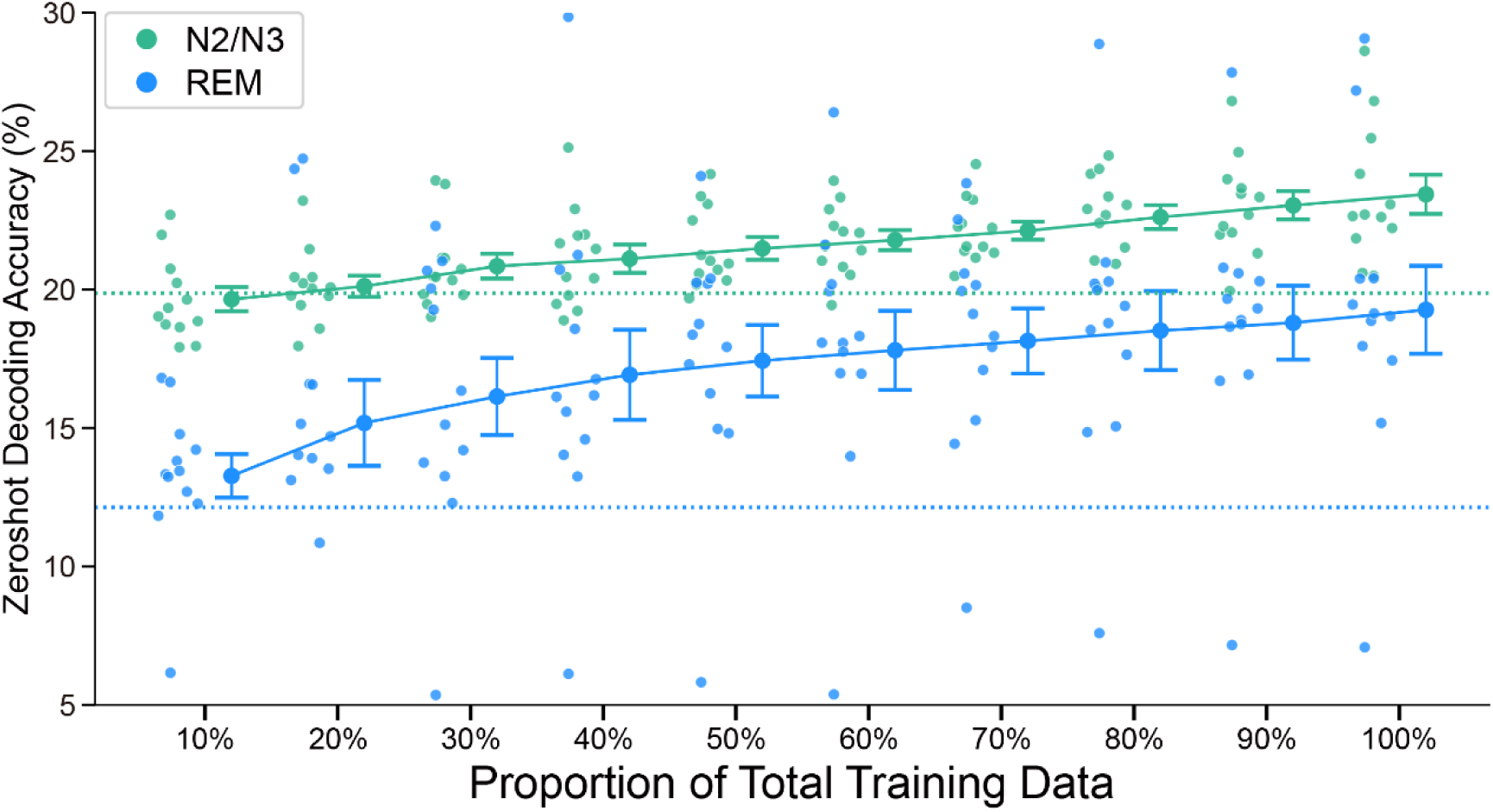
Zero-shot sleep decoding accuracy as a function of training data size. We evaluated the changes in zero-shot, cross-subject, sleep decoding performance under different training data volumes. This was done by randomly selecting 10% to 100% of the total training data to retrain the SI-MD model. The results for NREM2/3 stage are shown in green, and the results for REM stage are shown in blue. Each data point represents the result for a test subject.

**Extended Data Figure 3.**
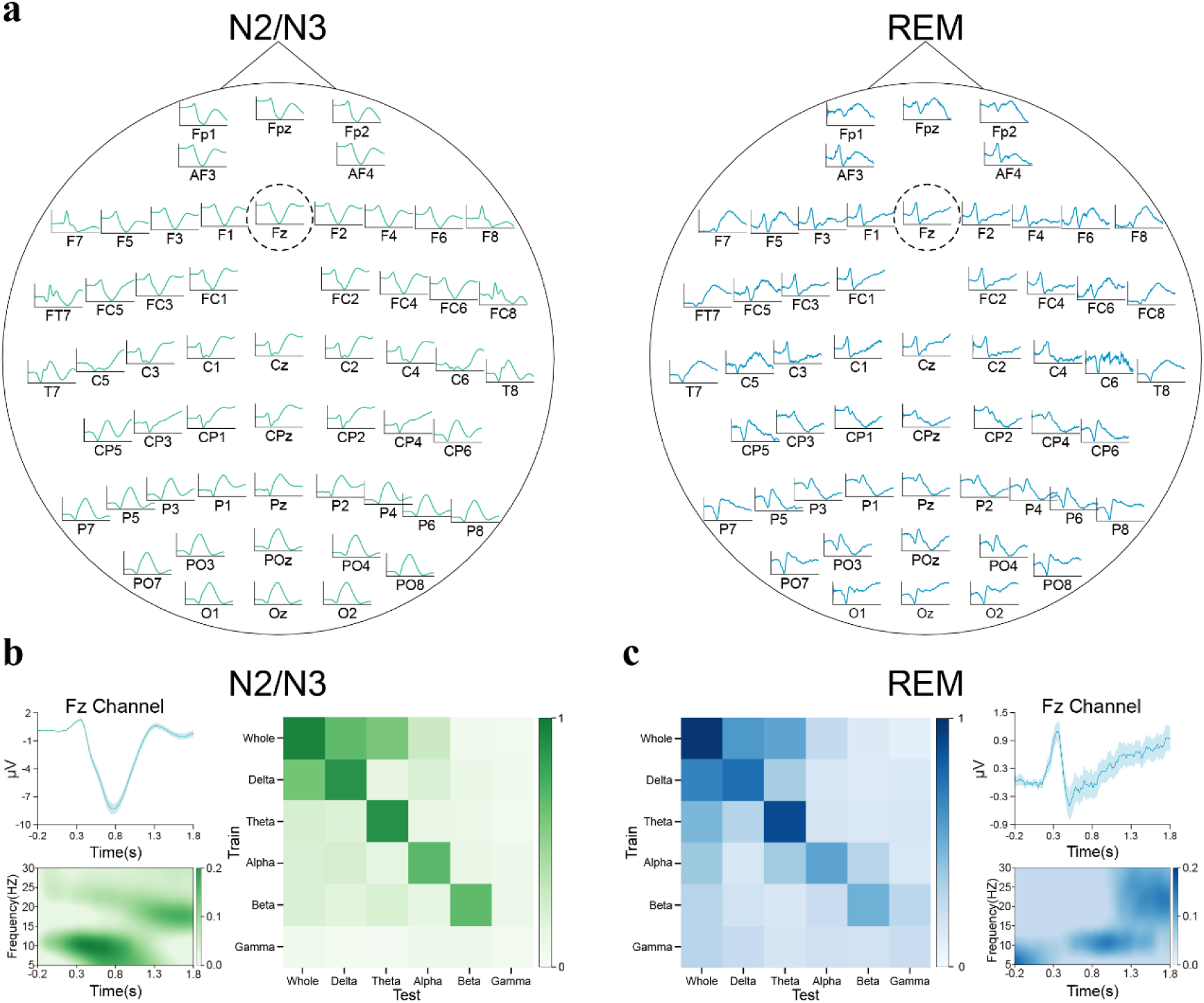
**ERPs and temporal frequency patterns for auditory-stimuli evoked neural response during sleep**. (**a**) Whole brain ERPs during sleep. ERPs of all EEG channels were shown separately for NREM2/3 (left panel) and REM (right panel) sleep stages. The time window spans from -0.2 to 1.8 seconds after stimulus onset. (**b**) Temporal frequency patterns of evoked neural response during NREM2/3 sleep. On the left, taking the Fz channel as an example, the upper left panel illustrates the ERP at the Fz channel during NREM2/3 stage, indicated in green, along with the corresponding standard errors across subjects. The lower left panel presents the time-frequency plots of the EEG signals from the Fz channel during this stage. The right panel shows the trained and tested results of the SI-MD based on specific frequency bands, with sleep decoding accuracy normalised between 0 and 1. (**c**) Temporal frequency patterns of evoked neural response during REM sleep, similar to panel (b), but for REM sleep, shown in blue.

**Extended Data Figure 4.**
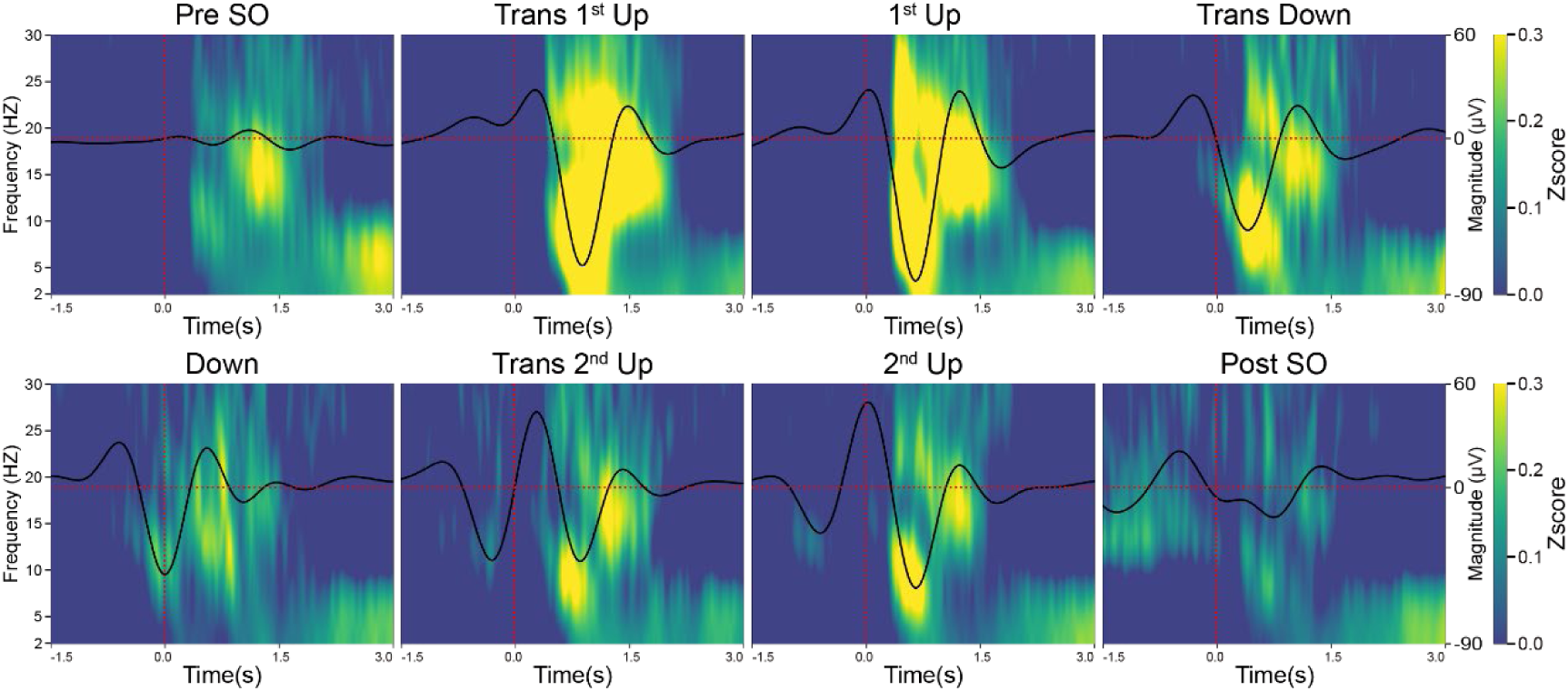
Temporal-frequency analysis of all eight SO-groups during NREM2/3 sleep. Temporal frequency analysis of EEG data from the Fz channel was shown separately for each of the eight SO-groups: ‘Pre SO’, ‘Transition to 1^st^ Up (Trans 1^st^ UP)’, ‘1^st^ Up’, ‘Transition to Down (Trans Down)’, ‘DOWN’, ‘Transition to 2^nd^ Up (Trans 2^nd^ UP)’, ‘2^nd^ Up’ and ‘Post SO’. The y-axis on the left side corresponds to the frequency range in the time-frequency plot, while the y-axis on the right corresponds to the magnitude in the ERP plot. For each SO-group, the time window is [-1.5s, 3.0s] relative to the sound stimulus onset. The vertical red lines mark the zero point in time, i.e., the onset of sound stimuli. The horizontal red line corresponds to the zero point of magnitude.

**Extended Data Figure 5.**
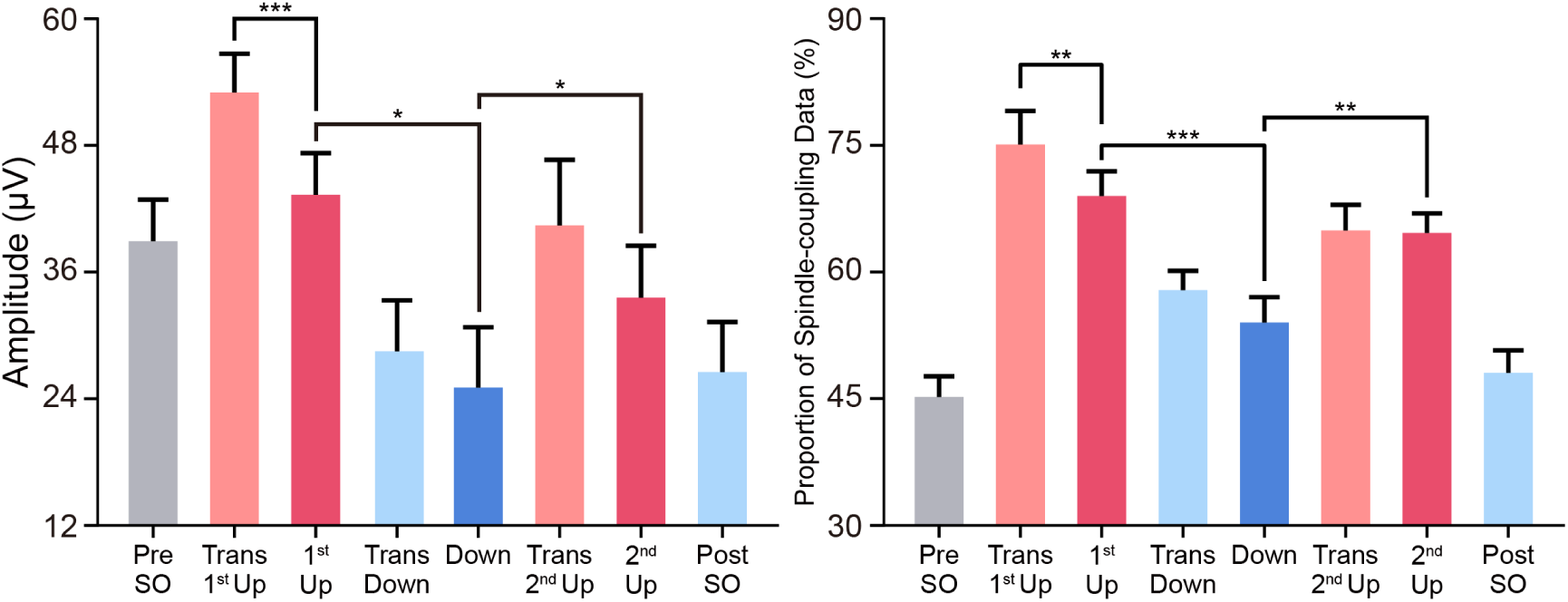
Distribution spindle events and mean magnitude across eight SO groups during NREM2/3 sleep. (**a**) The distribution of SO-spindle coupling in different SO groups, with the highest spindle coupling observed in the ‘Trans 1^st^ UP’ stage. (**b**) The distribution of the mean magnitude of ERPs in different SO groups, also showing the highest magnitude in the ‘Trans 1^st^ UP’ stage. The color code is in line with those used in Figure 5b. * *p*<0.05, ** *p*<0.01, *** *p*<0.001.

**Extended Data Figure 6.**
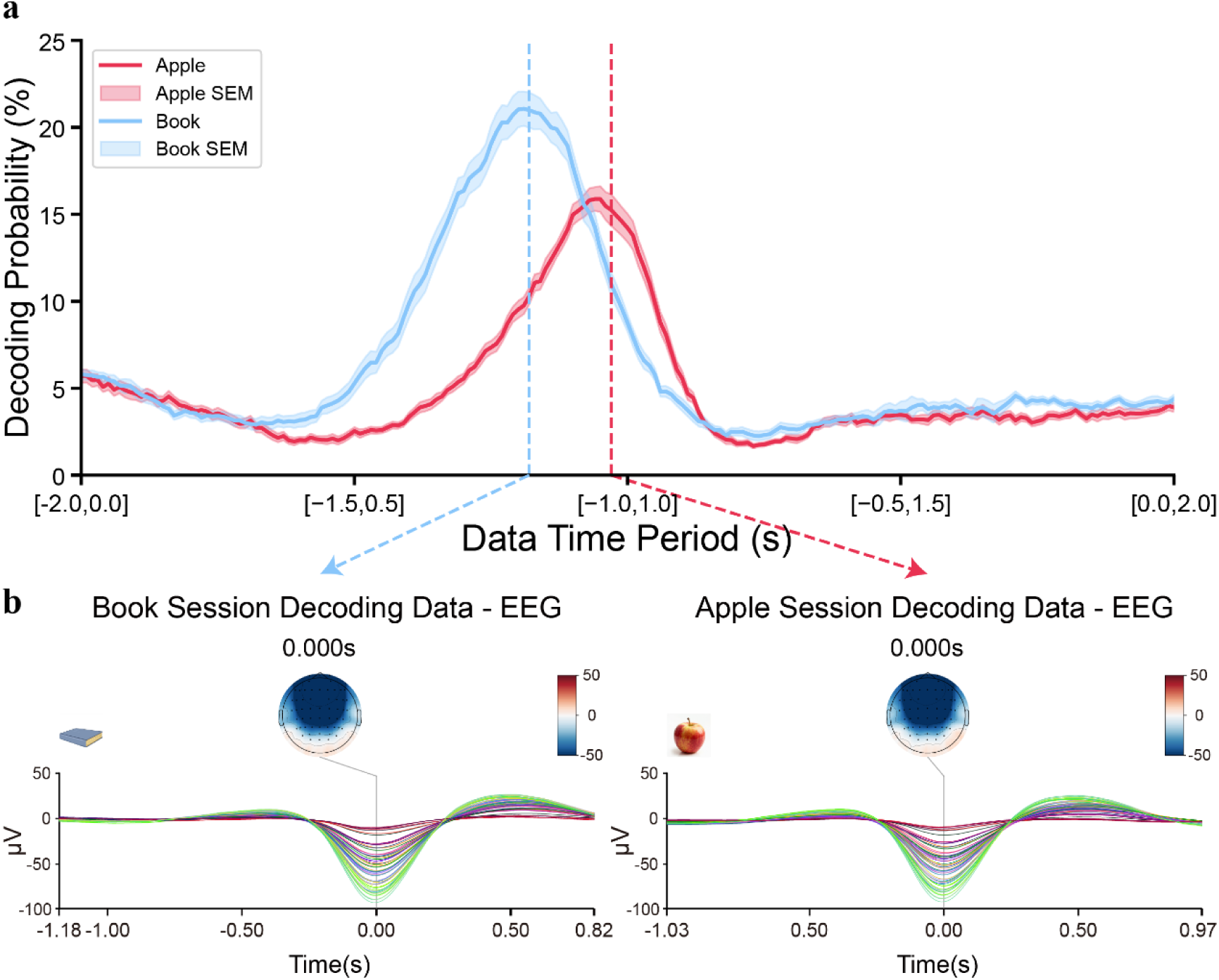
Decoding probability distributions and ERPs for the apple and book sessions in new experiment 1. (**a**) Mean decoding probability (± SE) in the apple session (red) and book session (blue) plotted against time relative to the slow-oscillation trough (time zero). The blue dashed vertical line indicates the optimal analysis window start in the book session, determined by splitting the data into validation and test sets and selecting the window with highest decoding probability on the validation set. The red dashed line marks the corresponding optimal window in the apple session, found by the same procedure. (**b**) Grand-average whole-brain ERPs and scalp topographies time-locked to the slow-oscillation trough for each session. ERPs span from 1.18 s before to 0.82 s after the trough in the book session and from 1.03 s before to 0.97 s after the trough in the apple session. Topographic maps are shown at the trough instant, with a common amplitude scale (–50 to 50 µV) to facilitate comparison of neural activation patterns.

**Extended Data Figure 7.**
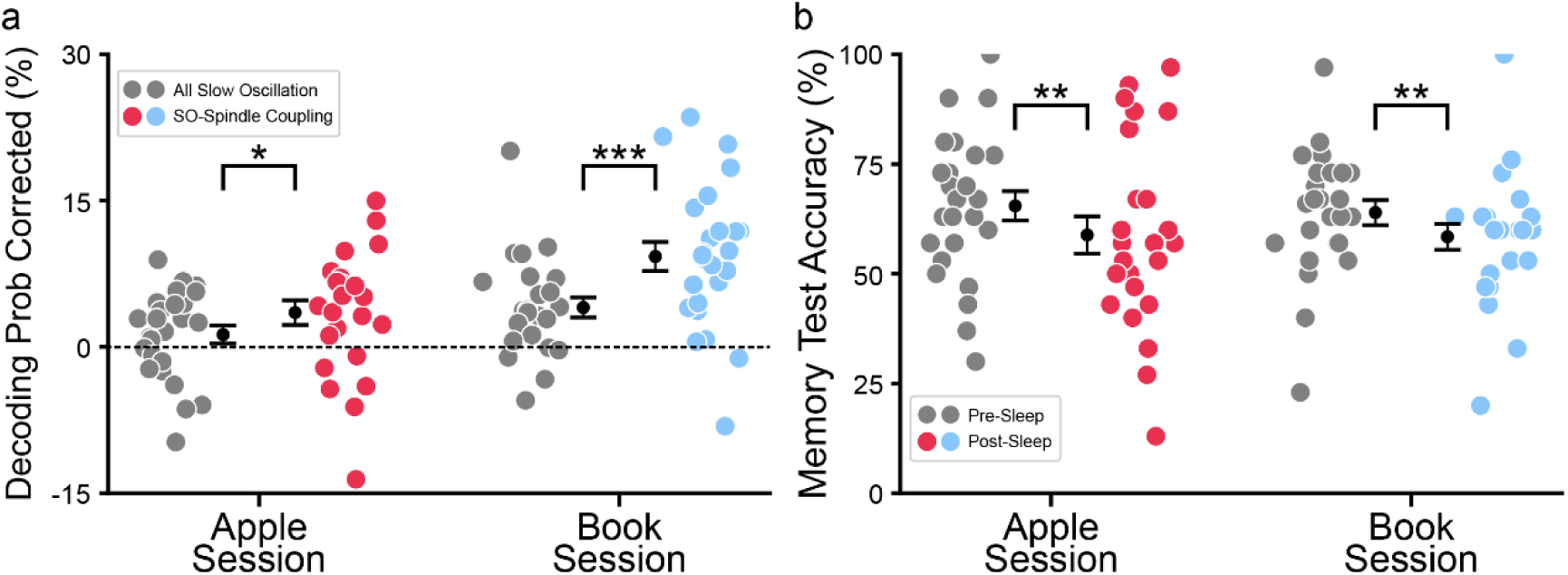
Decoding probabilities and behavioural memory performance before and after sleep in the new Experiment 1. (**a**) Decoding probabilities, corrected with reference data from the main-experiment, are shown for all SO segments (grey) and for SO-spindle coupled segments in the “apple” session (red) and “book” session (blue). Coupled segments yield the highest values. (**b**) Memory-test accuracy before sleep (grey) and after sleep for the “apple” session (red) and “book” session (blue). Error bars are mean ± SE. * *p*<0.05, ** *p*<0.01, *** *p*<0.001.

**Extended Data Figure 8.**
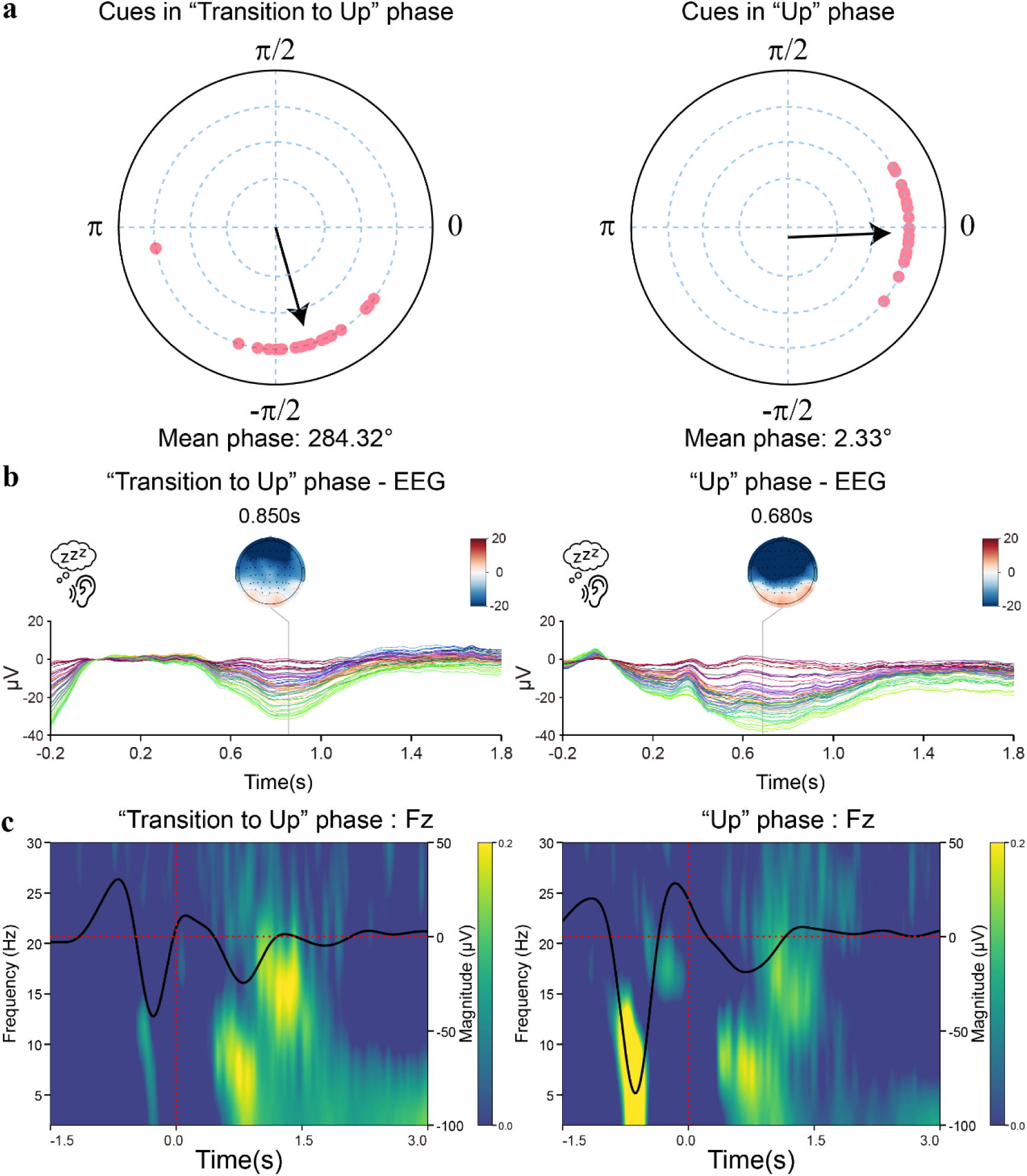
Phase distributions, ERPs and time-frequency representations for different sound-stimulation conditions in new experiment 2. (**a**) Circular histograms of cue delivery phases during targeted memory reactivation: “Transition to Up” (left) with mean phase 284.32° and “Up” (right) with mean phase 2.33°. Each dot represents one stimulation event; the solid line shows the mean resultant vector. (**b**) Grand-average ERPs from all EEG channels for cues delivered during the “Transition to Up” phase (left) and the “Up” phase (right), spanning -0.2 s to 1.8 s relative to stimulus onset. (**c**) Time-frequency plots from the Fz channel for the “Transition to Up” (left) and “Up” (right) phases, covering -1.5 s to 3.0 s around cue onset. The left y-axis indicates frequency (Hz) and the right y-axis shows ERP amplitude (µV). Vertical red lines denote cue onset (0 s) and the horizontal red line indicates zero amplitude.

**Extended Data Table 1.** Summary of sleep architecture metrics for the full cohort, training set, and test set.

| Sleep structure | All Participants | Train Set | Test Set |
| --- | --- | --- | --- |
| Sleep onset time(hh:mm) | 23:55 ± 00:03 | 23:55 ± 00:03 | 00:00 ± 00:07 |
| Geting up time (hh:mm) | 07:11 ± 00:02 | 07:10 ± 00:02 | 07:18 ± 00:06 |
| TST (min) | 355.77 ± 4.74 | 355.72 ± 5.10 | 356.33 ± 11.13 |
| WASO (min) | 52.91 ± 3.79 | 51.87 ± 4.04 | 63.67 ± 10.25 |
| SOL (min) | 13.90 ± 1.50 | 14.01 ± 1.61 | 12.71 ± 3.47 |
| SE (%) | 83.92 ± 1.01 | 84.07 ± 1.09 | 82.39 ± 2.31 |

Note. TST (Total Sleep Time): total minutes scored as “sleep” during time in bed; WASO: Wake After Sleep Onset; SOL: Sleep Onset Latency; SE (Sleep Efficiency): ratio of actual sleep time to total time in bed.

**Extended Data Table 2.** Number of presentations for each cue category during wakefulness, NREM sleep and REM sleep across all participants (Mean±SE).

| Number of Each category (unit) | Audio-awake | Image-awake | Audio-NREM | Audio-REM |
| --- | --- | --- | --- | --- |
| alarm | 143.59 ± 1.99 | 143.53 ± 1.95 | 93.30 ± 2.94 | 20.58 ± 1.08 |
| apple | 143.39 ± 1.92 | 143.33 ± 1.92 | 92.70 ± 2.93 | 20.79 ± 1.16 |
| ball | 143.79 ± 1.90 | 143.64 ± 1.97 | 94.77 ± 3.04 | 20.92 ± 1.10 |
| book | 143.46 ± 1.94 | 143.63 ± 1.91 | 93.02 ± 2.78 | 20.47 ± 1.16 |
| box | 143.48 ± 1.97 | 143.44 ± 1.95 | 93.60 ± 2.89 | 21.16 ± 1.13 |
| chair | 143.39 ± 1.92 | 143.53 ± 1.93 | 94.49 ± 3.12 | 20.24 ± 1.13 |
| kiwi | 143.38 ± 1.94 | 143.33 ± 1.90 | 93.28 ± 3.04 | 20.70 ± 1.02 |
| microphone | 143.47 ± 1.95 | 143.05 ± 1.99 | 92.66 ± 2.87 | 20.80 ± 1.10 |
| motorcycle | 143.56 ± 1.95 | 143.13 ± 2.02 | 93.13 ± 2.84 | 21.36 ± 1.15 |
| pepper | 143.09 ± 2.00 | 143.15 ± 1.99 | 94.95 ± 2.95 | 20.81 ± 1.14 |
| sheep | 143.81 ± 1.91 | 144.11 ± 1.90 | 93.26 ± 2.93 | 20.52 ± 1.14 |
| shoes | 143.75 ± 1.96 | 143.82 ± 1.93 | 93.50 ± 3.03 | 21.03 ± 1.17 |
| strawberry | 143.53 ± 1.95 | 143.59 ± 1.97 | 86.01 ± 3.23 | 19.21 ± 1.14 |
| tomato | 143.81 ± 1.97 | 143.78 ± 1.97 | 87.61 ± 3.18 | 19.46 ± 1.14 |
| watch | 143.93 ± 1.98 | 144.36 ± 1.96 | 86.88 ± 3.21 | 19.53 ± 1.15 |

Note. Audio-awake: Audio stimulus during Wakefulness; Image-awake: Image stimulus during Wakefulness; Audio-NREM: Audio stimulus during NREM sleep; Audio-REM: Audio stimulus during REM sleep.

**Extended Data Table 3.**
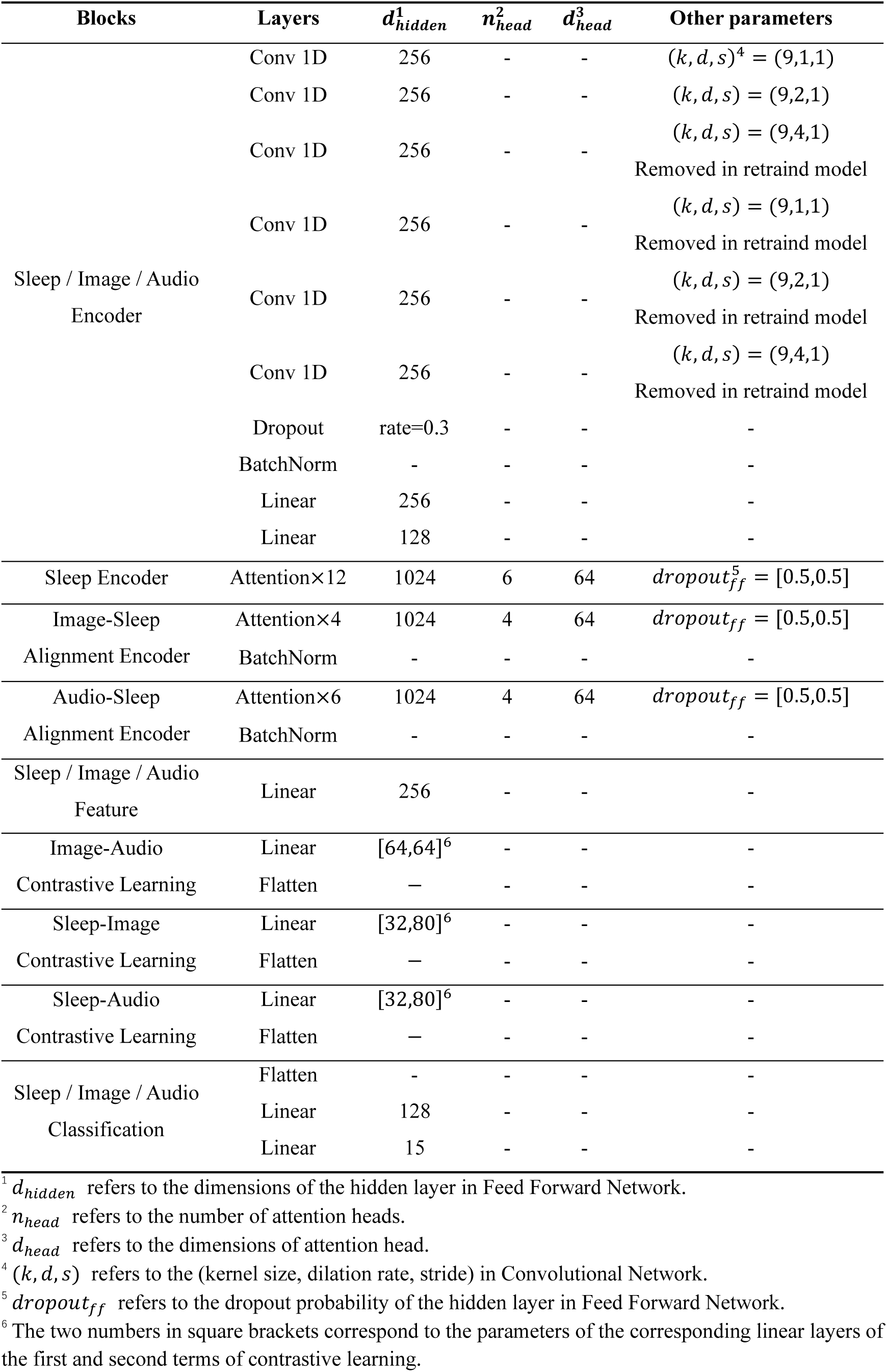
The configuration of SI-MD model.

**Extended Data Table 4.** Performance of different EEG-based ANN models in decoding NREM2/3 and REM sleep states. For each model, the 15-way classification accuracy (%) and the standard error (%) across test subjects are reported. The chance level is 1/15 = 6.7%.

| Models | Zero-shot |  | Finetune |  |
| --- | --- | --- | --- | --- |
|  | NREM 2/3 | REM | NREM 2/3 | REM |
| EEG-Net<sup>23</sup> | 17.30±0.37% | 14.05±1.11% | 20.03±0.66% | 21.38±1.95% |
| EEG-Conformer<sup>67</sup> | 19.12±0.48% | 14.81±0.79% | 23.74±0.83% | 24.61±1.82% |
| BrainBERT<sup>68</sup> | 12.50±0.78% | 10.59±1.31% | 17.30±1.44% | 18.21±2.02% |
| BIOT<sup>69</sup> | 16.72±0.52% | 13.15±1.08% | 22.80±1.19% | 23.93±2.09% |
| LaBram<sup>70</sup> | 17.91±1.61% | 13.64±2.11% | 21.99±2.14% | 22.27±2.92% |
| **SI-MD** | **22.82±0.52%** | **19.27±1.47%** | **30.81±0.65%** | **31.65±1.96%** |

**Extended Data Table 5.** Number of events in each SO phase category, reported for the full cohort as well as separately for the training and test sets (Mean±SE).

| Number of Each Phase | All Participants | Train Set | Test Set |
| --- | --- | --- | --- |
| Pre SO | 98.7 ± 3.52 | 98.33 ± 3.59 | 102.50 ± 15.39 |
| Trans 1st Up | 97.79 ± 4.33 | 98.37 ± 4.66 | 91.83 ± 9.36 |
| 1st Up | 59.35 ± 3.49 | 58.69 ± 3.70 | 66.08 ± 10.14 |
| Trans Down | 40.8 ± 1.67 | 41.28 ± 1.73 | 35.92 ± 6.38 |
| Down | 51.66 ± 2.05 | 52.71 ± 2.15 | 40.92 ± 6.09 |
| Trans 2nd Up | 40.39 ± 1.58 | 40.78 ± 1.62 | 36.33 ± 6.40 |
| 2nd Up | 69.38 ± 2.71 | 69.37 ± 2.70 | 69.5 ± 13.20 |
| Post SO | 61.17 ± 2.34 | 62.24 ± 2.41 | 50.25 ± 8.38 |

**Extended Data Table 6.** Sleep-related metrics for participants in Experiment 1 (no-TMR, spontaneous reactivation) and Experiment 2 (semantic-free cue TMR), presented as Mean±SE.

| Variables | New Experiment 1 | New Experiment 2 |
| --- | --- | --- |
| Sleep staging indices |  |  |
| Wake percent (%) | 24.68 ± 2.47 | 24.52 ± 3.66 |
| N1 percent (%) | 9.67 ± 0.93 | 8.71 ± 1.13 |
| N2 percent (%) | 35.14 ± 1.74 | 37.45 ± 2.96 |
| N3 percent (%) | 23.42 ± 2.09 | 21.84 ± 3.76 |
| REM percent (%) | 7.10 ± 1.47 | 7.48 ± 1.86 |
| TST (min) | 79.49 ± 3.31 | 80.20 ± 5.33 |
| SE (%) | 75.32 ± 2.47 | 75.48 ± 3.66 |
| Sleep rhythms |  |  |
| SO density (unit) | 6.47 ± 0.20 | 5.62 ± 0.32 |
| SO duration (s) | 1.42 ± 0.01 | 1.39 ± 0.01 |
| SO negative amplitudes (μV) | -49.74 ± 1.86 | -50.82 ± 2.42 |
| Spindle density (unit) | 9.88 ± 0.14 | 9.32 ± 0.24 |
| Spindle duration (s) | 0.89 ± 0.01 | 0.86 ± 0.02 |
| Spindle amplitudes (μV) | 12.65 ± 0.41 | 11.85 ± 0.54 |
| SO – spindle coupling density (unit) | 2.07 ± 0.07 | 1.70 ± 0.08 |
| SO – spindle coupling power (Z) | 0.77 ± 0.03 | 0.70 ± 0.05 |

**Note.** TST (Total Sleep Time): total minutes scored as “sleep” during time in bed; SE (Sleep Efficiency): ratio of actual sleep time to total time in bed.

**Extended Data Table 7.** Baseline decoding probabilities by category for SO-only and SO-spindle coupling epochs in stimulus-free sleep periods from the main experiment, time-locked to the same time intervals used in the new Experiment 1 (Mean ± SE).

| Decoding<br>Probabilities of<br>Each category<br>(SO Only / SO-spindle<br>Coupling) | Apple session<br>apple | Apple session<br>book | Book session<br>book | Book session<br>apple |
| --- | --- | --- | --- | --- |
| alarm | 0.92 ± 0.20 | 0.35 ± 0.10 | 0.53 ± 0.11 | 0.04 ± 0.02 |
| | 1.04 ± 0.29 | 0.30 ± 0.11 | 0.71 ± 0.16 | 0.03 ± 0.03 |
| apple | 14.05 ± 1.10 | 10.32 ± 0.57 | 14.09 ± 0.57 | 16.80 ± 1.28 |
| | 15.46 ± 1.22 | 10.80 ± 0.74 | 14.21 ± 0.73 | 18.22 ± 1.45 |
| ball | 3.97 ± 0.36 | 1.92 ± 0.16 | 1.16 ± 0.09 | 4.21 ± 0.51 |
| | 3.86 ± 0.37 | 1.84 ± 0.25 | 1.19 ± 0.15 | 3.86 ± 0.53 |
| book | 13.91 ± 0.88 | 18.59 ± 1.22 | 16.31 ± 0.93 | 17.42 ± 1.03 |
| | 15.31 ± 1.20 | 20.39 ± 1.68 | 17.42 ± 1.12 | 17.83 ± 1.10 |
| box | 14.12 ± 0.76 | 16.61 ± 0.98 | 20.9 ± 0.98 | 22.57 ± 1.39 |
| | 13.41 ± 0.88 | 16.62 ± 1.01 | 21.87 ± 1.16 | 21.96 ± 1.40 |
| chair | 3.33 ± 0.35 | 0.90 ± 0.14 | 1.80 ± 0.18 | 0.65 ± 0.17 |
| | 3.47 ± 0.49 | 0.89 ± 0.16 | 1.63 ± 0.16 | 0.83 ± 0.16 |
| kiwi | 1.59 ± 0.19 | 1.30 ± 0.18 | 1.50 ± 0.17 | 1.32 ± 0.16 |
| | 1.57 ± 0.19 | 1.28 ± 0.24 | 1.36 ± 0.23 | 1.28 ± 0.24 |
| microphone | 22.94 ± 1.03 | 24.48 ± 1.23 | 17.53 ± 1.15 | 15.69 ± 1.43 |
| | 21.22 ± 1.15 | 22.82 ± 1.36 | 16.47 ± 1.10 | 14.84 ± 1.55 |
| motorcycle | 7.85 ± 0.49 | 9.95 ± 0.78 | 11.02 ± 0.77 | 2.68 ± 0.46 |
| | 7.72 ± 0.59 | 9.38 ± 0.65 | 10.95 ± 0.70 | 2.58 ± 0.58 |
| pepper | 4.00 ± 0.34 | 3.90 ± 0.32 | 4.16 ± 0.33 | 0.25 ± 0.08 |
| | 3.59 ± 0.40 | 3.98 ± 0.38 | 3.62 ± 0.37 | 0.24 ± 0.11 |
| sheep | 1.99 ± 0.26 | 2.52 ± 0.20 | 2.63 ± 0.33 | 0.88 ± 0.15 |
| | 1.66 ± 0.25 | 2.43 ± 0.24 | 2.49 ± 0.41 | 0.65 ± 0.16 |
| shoes | 1.29 ± 0.12 | 2.79 ± 0.24 | 2.27 ± 0.23 | 1.53 ± 0.28 |
| | 1.34 ± 0.17 | 2.73 ± 0.32 | 2.28 ± 0.22 | 1.34 ± 0.29 |
| strawberry | 1.52 ± 0.13 | 0.79 ± 0.16 | 1.08 ± 0.13 | 0.12 ± 0.05 |
| | 1.42 ± 0.14 | 0.83 ± 0.17 | 1.01 ± 0.14 | 0.12 ± 0.05 |
| tomato | 3.28 ± 0.31 | 1.13 ± 0.19 | 1.76 ± 0.23 | 1.75 ± 0.25 |
| | 3.65 ± 0.50 | 1.13 ± 0.24 | 1.78 ± 0.31 | 1.99 ± 0.28 |
| watch | 5.24 ± 0.90 | 4.46 ± 0.63 | 3.26 ± 0.50 | 14.07 ± 1.21 |
| | 5.28 ± 1.13 | 4.60 ± 0.86 | 3.00 ± 0.39 | 14.24 ± 1.29 |

Note. Apple session apple / book: interval for decoding “apple” / “book” concept in “Apple” session of Experiment 1. Book session apple / book: interval for decoding “apple” / “book” concept in “Book” session of Experiment 1.

**Extended Data Video 1. Real-time sleep decoding demonstration.** We selected a representative segment from one test subject to illustrate the system’s operation. In the upper-left corner, ongoing sleep-stage classifications and decoding results are displayed, together with the top three class probabilities to guide the experimenter’s interpretation. On the right, the left image frame shows the cue-paired stimulus, while the right frame shows the corresponding image generated by the diffusion model based on the decoded class. The border of the generated-image panel turns green if the decoded class matches the cued category and red otherwise. Below these panels, the real-time EEG trace from the C3 channel (referenced to M2 and band-pass filtered 0.1-40 Hz) is plotted for the most recent 30 seconds, with all sound onsets marked by red vertical lines. At the very bottom, the current sleep stage for that 30-second epoch— determined by the SI-Staging model—is shown. The sleep staging updates every 3 seconds, whereas the SI-MD decoder runs only upon each sound cue, providing two decoding results per stimulus: one at 1.8 s (corresponding to the ERP window in Figure 3a) and a second at 3.5 s post-cue.

## References

1. Diekelmann, S. & Born, J. The memory function of sleep. Nat. Rev. Neurosci. 11, 114–126 (2010).

2. Rasch, B. & Born, J. About sleep’s role in memory. Physiol. Rev. (2013).

3. Klinzing, J.G., Niethard, N. & Born, J. Mechanisms of systems memory consolidation during sleep. Nat. Neurosci. 22, 1598–1610 (2019).

4. Brodt, S., Inostroza, M., Niethard, N. & Born, J. Sleep—A brain-state serving systems memory consolidation. Neuron 111, 1050–1075 (2023).

5. Gais, S. & Born, J. Declarative memory consolidation: mechanisms acting during human sleep. Learning & memory 11, 679–685 (2004).

6. Ngo, H.-V.V. & Staresina, B.P. Shaping overnight consolidation via slow-oscillation closed-loop targeted memory reactivation. Proc. Natl. Acad. Sci. USA 119, e2123428119. (2022).

7. Girardeau, G. & Lopes-dos-Santos, V. Brain neural patterns and the memory function of sleep. Science 374, 560–564 (2021).

8. Skaggs, W.E. & McNaughton, B.L. Replay of neuronal firing sequences in rat hippocampus during sleep following spatial experience. Science 271, 1870–1873. (1996).

9. Wilson, M.A. & McNaughton, B.L. Reactivation of hippocampal ensemble memories during sleep. Science 265, 676–679. (1994).

10. Staresina, B.P. Coupled sleep rhythms for memory consolidation. Trends Cognit. Sci. (2024).

11. Staresina, B.P., Niediek, J., Borger, V., Surges, R. & Mormann, F. How coupled slow oscillations, spindles and ripples coordinate neuronal processing and communication during human sleep. Nat. Neurosci. 26, 1429–1437 (2023).

12. Horikawa, T., Tamaki, M., Miyawaki, Y. & Kamitani, Y. Neural Decoding of Visual Imagery During Sleep. Science 340, 639–642 (2013).

13. Schönauer, M., et al. Decoding material-specific memory reprocessing during sleep in humans. Nat. Commun. 8, 15404 (2017).

14. Keefe, F.B., Johnson, L.C. & Hunter, E.J. EEG and autonomic response pattern during waking and sleep stages. Psychophysiology 8, 198–212 (1971).

15. Niedermeyer, E. The normal EEG of the waking adult. Electroencephalography: Basic principles, clinical applications, and related fields 167, 155–164 (2005).

16. Storch, C., Höhne, A., Holsboer, F. & Ohl, F. Activity patterns as a correlate for sleep–wake behaviour in mice. J. Neurosci. Methods 133, 173–179 (2004).

17. Sulaman, B.A., Wang, S., Tyan, J. & Eban-Rothschild, A. Neuro-orchestration of sleep and wakefulness. Nat. Neurosci. 26, 196–212 (2023).

18. Liu, Y., Nour, M.M., Schuck, N.W., Behrens, T.E.J. & Dolan, R.J. Decoding cognition from spontaneous neural activity. Nat. Rev. Neurosci. 23, 204–214. (2022).

19. Bishop, C.M. & Nasrabadi, N.M. Pattern recognition and machine learning (Springer, 2006).

20. Cortes, C. & Vapnik, V. Support-vector networks. Mach. Learn. 20, 273–297 (1995).

21. Cox, D.R. The Regression Analysis of Binary Sequences. Journal of the Royal Statistical Society: Series B (Methodological) 20, 215–232 (2018).

22. Cristianini, N. & Shawe-Taylor, J. An introduction to support vector machines and other kernel-based learning methods (Cambridge university press, 2000).

23. Lawhern, V.J., et al. EEGNet: a compact convolutional neural network for EEG-based brain--computer interfaces. J. Neural Eng. 15, 056013. (2018).

24. Ahmed, H., Wilbur, R.B., Bharadwaj, H.M. & Siskind, J.M. Object classification from randomized EEG trials. in Proceedings of the IEEE/CVF Conference on Computer Vision and Pattern Recognition 3845–3854 (2021).

25. Gifford, A.T., Dwivedi, K., Roig, G. & Cichy, R.M. A large and rich EEG dataset for modeling human visual object recognition. NeuroImage 264, 119754 (2022).

26. Kaneshiro, B., Perreau Guimaraes, M., Kim, H.-S., Norcia, A.M. & Suppes, P. A Representational Similarity Analysis of the Dynamics of Object Processing Using Single-Trial EEG Classification. PLoS One 10, e0135697 (2015).

27. Spampinato, C., et al. Deep learning human mind for automated visual classification. In Proceedings of the IEEE conference on computer vision and pattern recognition 6809–6817 (2017).

28. Chan, A.M., Halgren, E., Marinkovic, K. & Cash, S.S. Decoding word and category-specific spatiotemporal representations from MEG and EEG. Neuroimage 54, 3028–3039 (2011).

29. Frisby, S.L., Halai, A.D., Cox, C.R., Ralph, M.A.L. & Rogers, T.T. Decoding semantic representations in mind and brain. Trends Cognit. Sci. 27, 258–281 (2023).

30. Liu, H., Hajialigol, D., Antony, B., Han, A. & Wang, X. Eeg2text: Open vocabulary eeg-to-text decoding with eeg pre-training and multi-view transformer. arXiv preprint arXiv:2405.02165 (2024).

31. Wang, Z. & Ji, H. Open vocabulary electroencephalography-to-text decoding and zero-shot sentiment classification. in Proceedings of the AAAI Conference on Artificial Intelligence 5350–5358 (2022).

32. Tang, Z., Li, C. & Sun, S. Single-trial EEG classification of motor imagery using deep convolutional neural networks. Optik 130, 11–18 (2017).

33. Yang, L., Song, Y., Ma, K. & Xie, L. Motor imagery EEG decoding method based on a discriminative feature learning strategy. IEEE Trans. Neural Syst. Rehabil. Eng. 29, 368–379 (2021).

34. Al-Saegh, A., Dawwd, S.A. & Abdul-Jabbar, J.M. Deep learning for motor imagery EEG-based classification: A review. Biomed. Signal Process. Control 63, 102172 (2021).

35. Zhang, C., Kim, Y.-K. & Eskandarian, A. EEG-inception: an accurate and robust end-to-end neural network for EEG-based motor imagery classification. J. Neural Eng. 18, 046014 (2021).

36. Schirrmeister, R.T., et al. Deep learning with convolutional neural networks for EEG decoding and visualization. Hum. Brain Mapp. 38, 5391–5420 (2017).

37. Yu, Z., Chen, W. & Zhang, T. Motor imagery EEG classification algorithm based on improved lightweight feature fusion network. Biomed. Signal Process. Control 75, 103618 (2022).

38. Zhao, W., Jiang, X., Zhang, B., Xiao, S. & Weng, S. CTNet: a convolutional transformer network for EEG-based motor imagery classification. Sci. Rep. 14, 20237 (2024).

39. Schreiner, T., Petzka, M., Staudigl, T. & Staresina, B.P. Endogenous memory reactivation during sleep in humans is clocked by slow oscillation-spindle complexes. Nat. Commun. 12, 3112. (2021).

40. Hu, X., Cheng, L.Y., Chiu, M.H. & Paller, K.A. Promoting memory consolidation during sleep: A meta-analysis of targeted memory reactivation. Psychol. Bull. 146, 218 (2020).

41. Paller, K.A., Creery, J.D. & Schechtman, E. Memory and sleep: how sleep cognition can change the waking mind for the better. Annu. Rev. Psychol. 72, 123–150 (2021).

42. Rasch, B.r., Büchel, C., Gais, S. & Born, J. Odor cues during slow-wave sleep prompt declarative memory consolidation. Science 315, 1426–1429 (2007).

43. Radford, A., et al. Learning transferable visual models from natural language supervision. In International Conference on Machine Learning (ICML) 8748-8763 (PMLR, 2021).

44. McCarley, R.W. Neurobiology of REM and NREM sleep. Sleep medicine 8, 302–330 (2007).

45. Nielsen, T.A. A review of mentation in REM and NREM sleep:“covert” REM sleep as a possible reconciliation of two opposing models. Behav. Brain Sci. 23, 851–866 (2000).

46. Khosla, P., et al. Supervised contrastive learning. Advances in Neural Information Processing Systems (NeurIPS) 33, 18661–18673. (2020).

47. Chauvette, S., Volgushev, M. & Timofeev, I. Origin of active states in local neocortical networks during slow sleep oscillation. Cereb. Cortex 20, 2660–2674 (2010).

48. Lemieux, M., Chen, J.-Y., Lonjers, P., Bazhenov, M. & Timofeev, I. The impact of cortical deafferentation on the neocortical slow oscillation. J. Neurosci. 34, 5689–5703 (2014).

49. Volgushev, M., Chauvette, S., Mukovski, M. & Timofeev, I. Precise long-range synchronization of activity and silence in neocortical neurons during slow-wave sleep. J. Neurosci. 26, 5665–5672 (2006).

50. Hobson, J.A. REM sleep and dreaming: towards a theory of protoconsciousness. Nat. Rev. Neurosci. 10, 803–813 (2009).

51. Hobson, J.A., Stickgold, R. & Pace-Schott, E.F. The neuropsychology of REM sleep dreaming. Neuroreport 9, R1–R14 (1998).

52. Creery, J.D., Oudiette, D., Antony, J.W. & Paller, K.A. Targeted Memory Reactivation during Sleep Depends on Prior Learning. Sleep 38, 755–763 (2015).

53. Cairney, S.A., Lindsay, S., Sobczak, J.M., Paller, K.A. & Gaskell, M.G. The Benefits of Targeted Memory Reactivation for Consolidation in Sleep are Contingent on Memory Accuracy and Direct Cue-Memory Associations. Sleep 39, 1139–1150 (2016).

54. Fuentemilla, L., et al. Hippocampus-Dependent Strengthening of Targeted Memories via Reactivation during Sleep in Humans. Curr. Biol. 23, 1769–1775 (2013).

55. Xia, T., Antony, J.W., Paller, K.A. & Hu, X. Targeted memory reactivation during sleep influences social bias as a function of slow-oscillation phase and delta power. Psychophysiology 60, e14224 (2023).

56. Berry, R.B., et al. Rules for scoring respiratory events in sleep: update of the 2007 AASM manual for the scoring of sleep and associated events: deliberations of the sleep apnea definitions task force of the American Academy of Sleep Medicine. J. Clin. Sleep Med. 8, 597–619 (2012).

57. Iber, C. The AASM manual for the scoring of sleep and associated events: rules, terminology, and technical specification. (2007).

58. Vallat, R. & Walker, M.P. An open-source, high-performance tool for automated sleep staging. eLife 10, e70092 (2021).

59. Park, M.Y. & Hastie, T. L1-Regularization Path Algorithm for Generalized Linear Models. Journal of the Royal Statistical Society Series B: Statistical Methodology 69, 659–677 (2007).

60. Tibshirani, R. Regression Shrinkage and Selection via The Lasso: A Retrospective. Journal of the Royal Statistical Society Series B: Statistical Methodology 73, 273–282 (2011).

61. Tibshirani, R. Regression Shrinkage and Selection Via the Lasso. Journal of the Royal Statistical Society: Series B (Methodological) 58, 267–288 (2018).

62. Zou, H. & Hastie, T. Regularization and Variable Selection Via the Elastic Net. Journal of the Royal Statistical Society Series B: Statistical Methodology 67, 301–320 (2005).

63. Krizhevsky, A., Sutskever, I. & Hinton, G.E. ImageNet classification with deep convolutional neural networks. Commun. ACM 60, 84–90 (2017).

64. Vaswani, A., et al. Attention is all you need. Advances in Neural Information Processing Systems (NeurIPS) 30 (2017).

65. Dfossez, A., Caucheteux, C., Rapin, J., Kabeli, O. & King, J.-R. Decoding speech perception from non-invasive brain recordings. *Nat*. Mach. Intell. 5, 1097–1107 (2023).

66. Baevski, A., Zhou, Y., Mohamed, A. & Auli, M. wav2vec 2.0: A framework for self-supervised learning of speech representations. Advances in Neural Information Processing Systems (NeurIPS) 33, 12449–12460 (2020).

67. Song, Y., Zheng, Q., Liu, B. & Gao, X. EEG conformer: Convolutional transformer for EEG decoding and visualization. IEEE Trans. Neural Syst. Rehabil. Eng. 31, 710–719. (2022).

68. Wang, C., et al. BrainBERT: Self-supervised representation learning for intracranial recordings. arXiv preprint arXiv:2302.14367 (2023).

69. Yang, C., Westover, M. & Sun, J. Biot: Biosignal transformer for cross-data learning in the wild. Advances in Neural Information Processing Systems (NeurIPS) 36 (2024).

70. Jiang, W.-B., Zhao, L.-M. & Lu, B.-L. Large brain model for learning generic representations with tremendous EEG data in BCI. arXiv preprint arXiv:2405.18765 (2024).

71. Batterink, L.J., Creery, J.D. & Paller, K.A. Phase of spontaneous slow oscillations during sleep influences memory-related processing of auditory cues. J. Neurosci. 36, 1401–1409 (2016).

72. Hahn, M.A., Heib, D., Schabus, M., Hoedlmoser, K. & Helfrich, R.F. Slow oscillation-spindle coupling predicts enhanced memory formation from childhood to adolescence. elife 9, e53730 (2020).

73. Helfrich, R.F., Mander, B.A., Jagust, W.J., Knight, R.T. & Walker, M.P. Old brains come uncoupled in sleep: slow wave-spindle synchrony, brain atrophy, and forgetting. Neuron 97, 221–230. e224 (2018).

74. Muehlroth, B.E., et al. Precise slow oscillation–spindle coupling promotes memory consolidation in younger and older adults. Sci. Rep. 9, 1940 (2019).

75. Rombach, R., Blattmann, A., Lorenz, D., Esser, P. & Ommer, B. High-resolution image synthesis with latent diffusion models. in Proceedings of the IEEE/CVF conference on computer vision and pattern recognition 10684–10695 (2022).

76. Lawhern, V.J., et al. EEGNet: a compact convolutional neural network for EEG-based brain– computer interfaces. J. Neural Eng. 15, 056013 (2018).

77. Song, Y., Zheng, Q., Liu, B. & Gao, X. EEG conformer: Convolutional transformer for EEG decoding and visualization. IEEE Trans. Neural Syst. Rehabil. Eng. 31, 710–719 (2022).

78. Yang, C., Westover, M. & Sun, J. Biot: Biosignal transformer for cross-data learning in the wild. Advances in Neural Information Processing Systems (NeurIPS) 36, 78240–78260 (2023).

79. Albelwi, S. Survey on self-supervised learning: auxiliary pretext tasks and contrastive learning methods in imaging. Entropy 24, 551 (2022).

80. Jaiswal, A., Babu, A.R., Zadeh, M.Z., Banerjee, D. & Makedon, F. A survey on contrastive self-supervised learning. Technologies 9, 2 (2020).

81. Le-Khac, P.H., Healy, G. & Smeaton, A.F. Contrastive representation learning: A framework and review. IEEE Access 8, 193907–193934 (2020).

82. Liu, J., et al. Item-specific neural representations during human sleep support long-term memory. PLoS Biol. 21, e3002399 (2023).

83. Banville, H., Chehab, O., Hyvärinen, A., Engemann, D.-A. & Gramfort, A. Uncovering the structure of clinical EEG signals with self-supervised learning. J. Neural Eng. 18, 046020 (2021).

84. Thomas, A., Ré, C. & Poldrack, R. Self-supervised learning of brain dynamics from broad neuroimaging data. Advances in Neural Information Processing Systems (NeurIPS) 35, 21255–21269 (2022).

85. Grootswagers, T., Zhou, I., Robinson, A.K., Hebart, M.N. & Carlson, T.A. Human EEG recordings for 1,854 concepts presented in rapid serial visual presentation streams. Sci. Data 9, 3 (2022).

86. Hebart, M.N., et al. THINGS-data, a multimodal collection of large-scale datasets for investigating object representations in human brain and behavior. eLife 12, e82580 (2023).

87. Ackermann, S. & Rasch, B. Differential Effects of Non-REM and REM Sleep on Memory Consolidation? Curr. Behav. Neurosci. Rep. 14, 430 (2014).

88. Hobson, J.A., Pace-Schott, E.F. & Stickgold, R. Dreaming and the brain: Toward a cognitive neuroscience of conscious states. Behav. Brain Sci. 23, 793–842 (2000).

89. Walker, M.P., Liston, C., Hobson, J.A. & Stickgold, R. Cognitive flexibility across the sleep–wake cycle: REM-sleep enhancement of anagram problem solving. *Cogn*. Brain Res. 14, 317–324 (2002).

90. Schwartz, S., Clerget, A. & Perogamvros, L. Enhancing imagery rehearsal therapy for nightmares with targeted memory reactivation. Curr. Biol. 32, 4808–4816.e4804 (2022).

91. Siclari, F., et al. The neural correlates of dreaming. Nat. Neurosci. 20, 872–878 (2017).

92. Gramfort, A., et al. MEG and EEG data analysis with MNE-Python. Front. Neurosci. 7 (2013).

93. Li, A., Feitelberg, J., Saini, A.P., Höchenberger, R. & Scheltienne, M. MNE-ICALabel: Automatically annotating ICA components with ICLabel in Python. J. Open Source Softw. 7, 4484 (2022).

94. Ioffe, S. & Szegedy, C. Batch normalization: Accelerating deep network training by reducing internal covariate shift. in International Conference on Machine Learning (ICML) 448-456. (PMLR, 2015).

95. Gupta, N., Smith, J., Adlam, B. & Mariet, Z.E. Ensembles of classifiers: a bias-variance perspective. Transactions on Machine Learning Research (2022).

96. Mohammed, A. & Kora, R. A comprehensive review on ensemble deep learning: Opportunities and challenges. Journal of King Saud University - Computer and Information Sciences 35, 757–774 (2023).

97. Staresina, B.P., et al. Hierarchical nesting of slow oscillations, spindles and ripples in the human hippocampus during sleep. Nat. Neurosci. 18, 1679–1686. (2015).

